# A high-throughput compound screen identifies multiple druggable targets in Plasmodium falciparum transmission stages

**DOI:** 10.64898/2026.09.10.747198

**Authors:** Leonie Seefeldt, Christin Gumpp, Olivia Carril, Jerome Eberhardt, Sylwia Boltryk, Armin Passecker, Basil Tino Thommen, Kleopatra Sifoniou, Christian Scheurer, Christoph Fischli, Deszö Imre Babai, Amr Hamed Mahmoud, Leila Tamara Alexander, Steffen Renner, W. Armand Guiguemde, Luying Pei, Christof Grüring, Heike Butendeich, Daniela Siebert, Judith Straimer, Victor Tobiasson, Sebastian Baumgarten, Markus Alexander Lill, Daniel Baeschlin, Matthias Rottmann, Till Steffen Voss, Nicolas Michel Beat Brancucci

## Abstract

Most antimalarials are ineffective against the sexual transmission stages, known as gametocytes, of the malaria parasite *Plasmodium falciparum.* Their low sensitivity to drugs is attributed to limited compound uptake and a poorly understood form of cellular quiescence. Our current understanding of druggable transmission-blocking processes is therefore limited. Based on genetically engineered parasites that facilitate the mass production of synchronous mature gametocytes, we developed a high throughput drug screening platform that allowed us to test more than 50,000 compounds for gametocytocidal effects in one day. By screening a diversity-oriented library, we identified over 40 molecules that kill mature gametocytes in the low nanomolar range. Using resistance selection coupled to whole genome sequencing and drug-target interaction modelling, we followed up on three chemically tractable compounds that are also highly active against asexual parasites and prevent gametocyte transmission to mosquitoes. We show that the compound ONX-0914, a specific inhibitor of the β5i/LMP7 subunit of human immunoproteasomes, targets the parasite proteasomal β5 subunit. In contrast, the compounds CR-1-31-B and brusatol interfere with translation by targeting eukaryotic initiation factor 4A (eIF4A) and the peptidyl transferase center (PTC) of the 80S ribosome, respectively. Interestingly, parasite resistance to brusatol, a broad-spectrum antitumor drug, is linked to the differential modification of specific rRNA bases near the ribosomal A-site, mediated by altered base specificity of a rRNA methyltransferase. In summary, we successfully combined high-throughput compound screening with drug target deconvolution to reveal the targets and mode-of-action for three potent gametocytocidal molecules and discover the mechanism of resistance to the anti-tumorigenic drug brusatol. In addition to critically advancing our understanding of mature gametocyte biology and druggable processes in *P. falciparum* transmission stages, our observations made with brusatol-resistant parasites may become relevant for anti-cancer drug research.

**One sentence summary:** We identified three malaria transmission-blocking drug targets by combining high-throughput compound screening with in vitro resistance selection, whole-genome sequencing and drug target deconvolution.

## INTRODUCTION

Malaria continues to threaten public health in many countries worldwide. Despite the progress made toward eliminating the disease in some regions, 263 million clinical malaria cases and close to 610,000 deaths were reported for 2024, of which two-thirds occurred in eleven African countries (World Health Organization, 2025). Effective transmission-blocking interventions are critical for maintaining low infection rates and achieving current and future elimination goals, but they are complicated by the fact that gametocytes, the parasite stages responsible for malaria transmission, are insensitive to most antimalarial drugs. Furthermore, parasites partially resistant to the frontline artemisinin-based combination therapies (ACTs) continue to spread and further endanger disease intervention efforts (Dondorp *et al*., 2011; Rabinovich *et al*., 2017; Dhorda, Amaratunga and Dondorp, 2021).

*Plasmodium falciparum* is the most virulent of the six *Plasmodium* species infecting humans and responsible for the majority of malaria-related morbidity and mortality. Following infection and intra-hepatocytic replication, parasites start the symptomatic phase of infection by entering the bloodstream, where they undergo continuous cycles of red blood cell (RBC) invasion, intra-erythrocytic replication via schizogony, and the release of daughter merozoites that invade new RBCs. During each of these asexual replication cycles, a small proportion of parasites commit to the formation of gametocytes, the only stage capable of infecting the mosquito vector. Since these sexual transmission stages are non-replicating, parasites have to balance investments into chronic infection (asexual replication) versus transmission (gametocyte formation) (Abdi, 2023; Sollelis, Howick and Marti, 2024). Gametocytes are thus produced at variable but generally low rates, and the parasites evolved sophisticated mechanisms to control the switch between asexual replication and gametocyte formation (Voss and Brancucci, 2024). Specifically, parasites use environmental cues to regulate transcriptional silencing or activation of the gene encoding PfAP2-G – the master transcription factor necessary and sufficient to initiate sexual commitment and subsequent gametocyte differentiation (Kafsack, 2014; Sinha *et al*., 2014). Typically, the *pfap2-g* locus is epigenetically silenced by mechanisms dependent on histone deacetylase 2 (PfHDA2) and histone 3 lysine 9 trimethylation (H3K9me3)/heterochromatin protein 1 (PfHP1), thereby preventing excessive investments into gametocyte production (Brancucci *et al*., 2014; Coleman *et al*., 2014). In response to environmental stressors, including limiting levels of substrates required for lipid biogenesis (Pessi, Kociubinski and Mamoun, 2004; Brancucci *et al*., 2017; Wein *et al*., 2018), parasites induce expression of gametocyte development 1 (PfGDV1), a nuclear factor that interacts with and expels PfHP1 from the *pfap2-g* locus and thereby triggers PfAP2-G expression (Filarsky *et al*., 2018). Using these interlinked control layers, *P. falciparum* parasites are capable of adapting gametocyte production rates under changing environmental conditions (Voss and Brancucci, 2024).

Sexual commitment is the first known step in the process of gametocytogenesis and takes place in the intra-erythrocytic cycle preceding gametocyte differentiation; PfAP2-G-expressing schizonts produce sexual ring stage progeny that will develop into gametocytes rather than undergoing another round of asexual replication (Bruce *et al*., 1990; Voss and Brancucci, 2024). During sexual differentiation, gametocytes sequester away from peripheral circulation and accumulate primarily within the bone marrow, where they develop through four different morphological stages (stages I-IV) before reaching maturity after 10 to 12 days (Venugopal, 2020). Mature stage V gametocytes adopt an eponymous falciform (crescent) shape and re-enter the bloodstream, circulating for days to weeks to maximize transmission success (Bousema *et al*., 2010; Venugopal *et al*., 2020). During this phase, stage V gametocytes exist in a quiescent state and are believed to wind down cellular processes at multiple levels, including translation and metabolism (Keroack and Duraisingh, 2022). When ingested by a female *Anopheles* spp. mosquito, stage V gametocytes egress from their host cell and form female and male gametes that undergo fertilization and transform into a motile ookinete capable of crossing the midgut epithelium. Here, oocyst development results in the release of thousands of sporozoites that eventually migrate to the salivary glands ready to infect another human host.

The distinct developmental fates of macrogametes (females) and microgametes (males) are already predetermined during gametocyte differentiation, as evident from their sex-specific transcriptional and proteomic signatures (Lasonder *et al*., 2016; Miao *et al*., 2017; Jeninga *et al*., 2023; Dogga *et al*., 2024; Mohammed, 2024). Female gametocytes transcribe and store mRNA from numerous genes in translationally repressed ribonucleoprotein complexes containing DOZI (Development Of Zygote Inhibited) (Mair *et al*., 2006, 2010; Guerreiro *et al*., 2014; Min *et al*., 2024). By contrast, male gametocytes prepare for rapid genome replication, endomitosis and cytokinesis required for the release of eight microgametes (exflagellation). In the mosquito midgut, gametocytes activate within seconds in response to a drop in temperature and exposure to xanthurenic acid (Billker *et al*., 1998; Garcia *et al*., 1998; Arai *et al*., 2001). Orchestrated by a receptor complex containing a guanylyl-cyclase, gametocyte activation is initiated by the cyclic GMP-mediated modulation of protein kinase G (PfPKG) activity and the subsequent increase in cellular calcium levels (Muhia *et al*., 2001; McRobert *et al*., 2008; Brochet and Billker, 2016). The consequent activation of downstream effectors results in the formation of male and female gametes (Janse *et al*., 1988; Fang *et al*., 2017; Invergo *et al*., 2017; Matthews *et al*., 2022). After fertilization, the release of maternal transcripts from DOZI-mediated repression is important for zygote formation and further parasite development within the mosquito (Mair, 2006; Mair *et al*., 2010; Guerreiro *et al*., 2014).

Mature stage V gametocytes use several regulatory layers to enter a quiescent developmental state, the mechanisms of which are still rather poorly understood (Keroack and Duraisingh, 2022). Translational control through protein kinase 4 (PfPK4)-mediated phosphorylation of eukaryotic initiation factor 2ɑ (PfeIF2ɑ) (Zhang *et al*., 2012) as well as changes in metabolic activity (Sinden and Smalley, 1979; Delves *et al*., 2013; Lamour *et al*., 2014; Gulati *et al*., 2015) have been reported. More recently, Grünebast and colleagues showed that ribosomal RNA (rRNA) is gradually degraded from stage II gametocytes onwards, possibly resulting in reduced translational activity in mature gametocytes (Grünebast *et al*., 2025). Furthermore, changes in membrane composition during gametocyte maturation are linked to reduced compound uptake (Tran *et al*., 2016; Ridgway *et al*., 2022; Naude *et al*., 2024). These and other unknown mechanisms likely contribute in a major way to the general insensitivity of mature gametocytes to drug-mediated killing.

Identifying chemical compounds active against stage V gametocytes holds promise not only for discovering new transmission-blocking drug candidates and druggable targets, but also for generating new insight into the biology of these quiescent cells. Because gametocytes become increasingly tolerant to drugs during their maturation process, drug screening efforts ideally focus on mature stage V gametocytes. However, the *in vitro* production of these stages is complicated by the generally low rate at which parasites commit to the sexual pathway and differentiate into gametocytes (Voss and Brancucci, 2024). Despite this limitation, various screening efforts, including high-throughput-compatible approaches, have successfully been developed and applied to identify molecules with potent activity against late stage gametocytes (Birkholtz, Alano and Leroy, 2022; Paonessa *et al*., 2022; Reader, Van Der Watt and Birkholtz, 2022; Van Der Watt, Reader and Birkholtz, 2022; Brancucci *et al*., 2025). To generate gametocyte populations suitable for drug screening, most methods exploit the fact that sexual conversion rates increase in response to environmental stressors, including nutrient restriction caused by high parasitemia and/or the addition of parasite-conditioned/spent medium (Williams, 1999; Fivelman *et al*., 2007; Delves *et al*., 2016). While these methods allow for increased gametocyte yields, the resulting populations often contain stage V gametocytes mixed with earlier stages, limiting the confidence with which transmission-blocking compounds can reliably be identified. The recent development of transgenic NF54/iGP parasite lines for the mass production of synchronous stage V gametocytes via conditional overexpression of PfGDV1 mitigated these limitations and offered new avenues for transmission-blocking drug discovery and development (Boltryk *et al*., 2021; Brancucci *et al*., 2025; Schäfer *et al*., 2026). For instance, we recently engineered the NF54/iGP1_RE9H^ulg8^ line that expresses the red-shifted firefly luciferase RE9H controlled by the gametocyte-specific *ulg8* promoter (Siciliano *et al*., 2017; Boltryk *et al*., 2021; Brancucci *et al*., 2025). Using this cell line, we established an improved *in vitro* assay for stage V gametocytocidal drug discovery using small-to medium-sized chemical libraries as well as a humanized mouse model for gametocyte infection facilitating the pre-clinical efficacy testing of transmission-blocking drug candidates *in vivo* (Brancucci *et al*., 2025).

Here, we engineered the NF54/iGP1_RE9H^pfs16^ reporter line carrying a RE9H expression cassette driven by the strong gametocyte-specific *pfs16* promoter (Dechering *et al*., 1999) and developed a highly sensitive and robust *in vitro* assay for identifying compounds active against stage V gametocytes at a throughput of more than 50,000 compounds per day. We screened a diversity-oriented compound library consisting of 51,185 molecules under high-throughput conditions and identified potent gametocytocidal compounds that prevent mosquito infection in the low nanomolar range. By coupling *in vitro* resistance selection to whole genome sequencing and drug-target interaction modelling, we revealed the molecular targets, modes of action and resistance mechanisms for three of the most active hits, ONX-0914, CR-1-31-B and brusatol. Together, these efforts identified several highly potent malaria transmission-blocking compounds by high-throughput screening and indicated that stage V gametocytes are particularly vulnerable to compounds targeting proteostasis.

## RESULTS

### Engineering of the NF54/iGP1_RE9H^pfs16^ parasite line for gametocytocidal drug screening

To be able to screen for compounds active against stage V gametocytes at high throughput, we made use of the inducible gametocyte producer line NF54/iGP1 (Boltryk *et al*., 2021). These parasites carry a chromosomal expression cassette, inserted into the non-essential *cg6* (*glp3*) locus (PF3D7_0709200), encoding GDV1 fused to green fluorescent protein (GFP) and the FKBP destabilization domain (DD), followed by a *glmS* ribozyme sequence element (GDV1-GFP-DD-*glmS*). When cultured in the presence of glucosamine (GlcN; activates *glmS* ribozyme activity in the 3’ untranslated region) and the absence of the DD-stabilizing compound Shield-1 (Armstrong and Goldberg, 2007; Parichat Prommana *et al*., 2013), expression of the GDV1 fusion protein is prevented. However, upon addition of Shield-1 and withdrawal of GlcN, GDV1-GFP-DD is stably expressed and induces sexual commitment in a large fraction of parasites, resulting in the mass production of highly synchronous gametocytes (Boltryk *et al*., 2021). Here, we further modified NF54/iGP1 parasites by inserting directly downstream of the ectopic *gdv1-gfp-dd-glmS* locus a second expression cassette encoding the red-shifted ATP-dependent firefly luciferase RE9H driven by the gametocyte-specific *pfs16* promoter (Dechering *et al*., 1999) (NF54/iGP1_RE9H^pfs16^) (Fig. 1A and Fig. S1). As shown in Fig. 1B, luminescence intensity was linearly correlated with the number of viable gametocytes over a wide range and sensitive detection with robust signal above background was obtained for less than 100 gametocytes per well.

We next tested the suitability of NF54/iGP1_RE9H^pfs16^ gametocytes for drug screening by following the protocol recently developed for the NF54/iGP1_RE9H^ulg8^ reporter line (Brancucci *et al*., 2025). In brief, sexual commitment was induced by removing GlcN and adding 675 nM Shield-1 (+Shield/-GlcN) to synchronized asexual parasites (0-8 hours post invasion (hpi), 2% parasitemia). After 48 hours, the ring stage progeny (containing a high proportion of sexual ring stages) was kept in regular culture medium supplemented with 50 mM N-acetylglucosamine (GlcNAc) during days 1 to 6 of gametocyte development to selectively eliminate asexual parasites (Ponnudurai *et al*., 1986; Fivelman *et al*., 2007). On day 12, cultures typically contained pure stage V gametocytes at a gametocytemia above 3%. At this point, gametocytemia and hematocrit were adjusted to 0.3% and 1%, respectively, and the gametocytes were transferred to wells of different plate formats (100 µl/well for the 96-well format; 7.5 µl/well for the 1536-well format) pre-loaded with test compounds (Fig. 1C). To validate the high-throughput setup, we tested the effect of gold standard compounds known to be active against stage V gametocytes (methylene blue, MB) or inactive (chloroquine, CQ). Luminescence was recorded after an incubation period of 72 hours, revealing that MB and CQ elicited the expected dose-response in both the 96-well and 1536-well assay formats (Fig. 1D).

**Figure 1.**
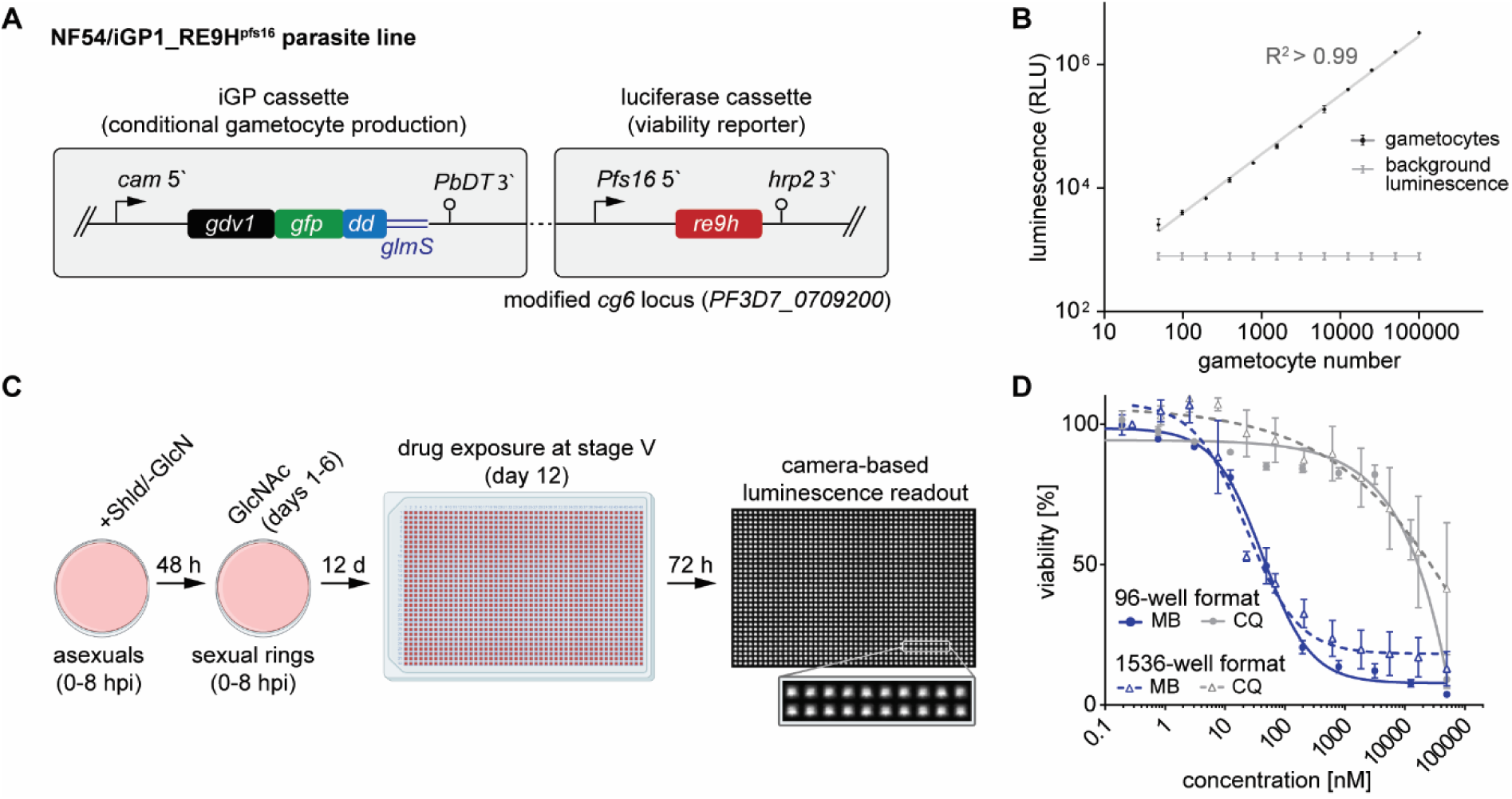
The NF54/iGP1_RE9H^pfs16^ line allows probing gametocyte viability under high throughput screening (HTS) conditions. **(A)** Schematic map of the inducible gametocyte producer (iGP) and *re9h* viability reporter cassettes inserted into the non-essential *glp3/cg6* locus in NF54/iGP1_RE9H^pfs16^ parasites. **(B)** Luminescence intensity linearly correlates with the number of viable stage V gametocytes across a large range. **(C)** Experimental setup of the 1536-well gametocytocidal drug activity assay. **(D)** Comparison of dose-dependent MB and CQ activity against NF54/iGP1_RE9H^pfs16^ stage V gametocytes (day 12) quantified using the 96-well and 1536-well formats. Values on the y-axis represent luminescence normalized to the mean signal emitted from untreated cells. Mean and standard error of the mean (s.e.m) of three biological replicates (n=3) are shown.

### High-throughput screening of a chemically diverse compound library identifies potent stage V gametocytocidal compounds

Next, we screened a large Novartis compound library for molecules able to kill stage V gametocytes using the 1536-well plate format. This library consisted of different sets of non-proprietary compounds, including molecules with known mammalian targets (MoA, mode of action library, 841 compounds (Canham *et al*., 2020), natural products (NAPR, 2475 compounds) as well as a diversity-oriented collection of molecules (Schuffenhauer *et al*., 2020), totaling 51,185 unique compounds (Dataset S1). Using an automated dispenser (AquaMax DW4 microplate dispenser, Molecular Devices), 7.5 µl of gametocyte suspension (0.3% gametocytemia, 1% hematocrit; approx. 2,000 gametocytes/well) were added to each well of the plates pre-spotted with 7.5 nl of 5 mM compound (in 90% dimethyl sulfoxide; DMSO) by an acoustic liquid handler (Echo 550, Beckman Coulter). In this setup, less than 1 µl of a standard NF54/iGP1_RE9H^pfs16^ stage V gametocyte culture (day 12) is sufficient per test condition, which allowed screening all 51,185 compounds on a single day using 60 ml of gametocyte culture (∼3% gametocytemia, 3% hematocrit). Immediately after dispensing, gametocytes were incubated for 72 hours at a drug concentration of 5 µM. Gametocyte viability was subsequently assessed by adding the substrate D-Luciferin (2.5 µl at 1 µg/µl, Perkin Elmer) and reading out luminescence intensity using a camera-based plate reader (Fig. 2A). Wells containing different concentrations of MB (800 nM, 10 µM and 50 µM) and the compound vehicle (DMSO at 0.05% final concentration) were used as treated and untreated controls, respectively. Gametocytocidal effects of the test compounds (reduction in gametocyte viability) were quantified by normalizing the raw luminescence counts from each well to the mean signals obtained from the negative controls (untreated; 0% activity) and positive controls (50 µM MB-treated, -100% activity). Assay quality was monitored on a plate-by-plate basis with Z’ factors consistently remaining above 0.75 (0.82 ± 0.05 SD) (Fig. 2B).

**Figure 2.**
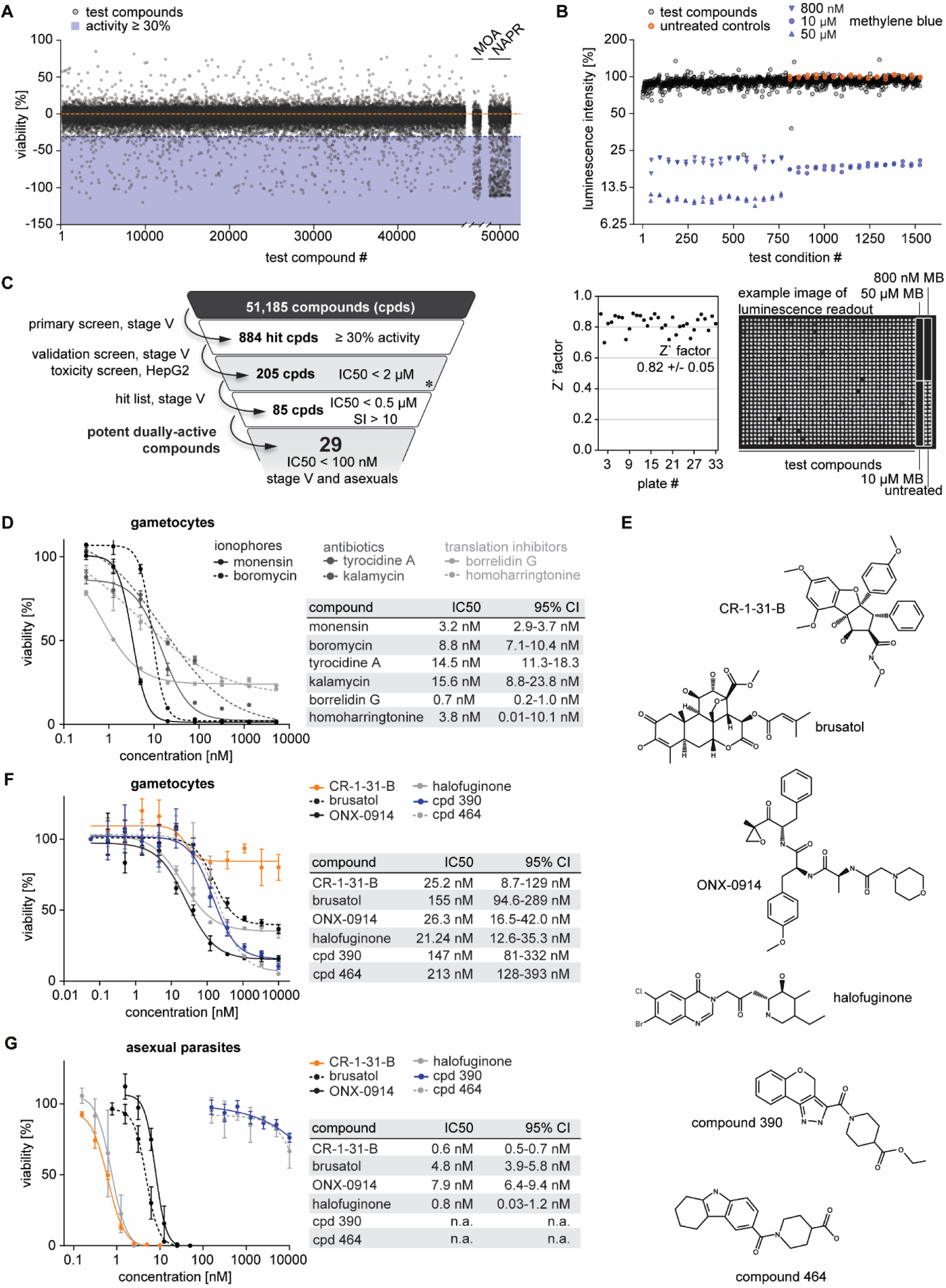
Identification of stage V gametocytocidal compounds via high throughput screening. **(A)** Primary screening of 51,185 compounds. The 30% activity threshold is highlighted by the blue-shaded area (left panel). NAPR, natural product library; MOA, mode of action library. **(B)** Results of an example plate with untreated and treated controls shown in orange and blue, respectively (top). Z’ factors calculated for each screening plate are shown below. Scores were calculated using untreated gametocytes and gametocytes treated with 50 µM MB. Means and standard deviations are indicated. An example image of the luminescence-based viability readout is shown on the right. **(C)** Hit triaging. Inclusion criteria and number of compounds fulfilling the respective criteria are indicated. *asexual stage IC_50_ data is available for 156 of the 205 compounds with gametocytocidal activity < 2 µM. **(D)** Dose-response activity of selected hit compounds on stage V gametocytes (day 12) determined in the validation screen [mean ± s.e.m., n=1 (technical quadruplicates in the 1536-well assay format)]. IC_50_ values and 95% confidence intervals (95% CI) are indicated. **(E)** Chemical structures of selected hit compounds. **(F)** Dose-response activity of selected hit compounds on stage V gametocytes performed using the 96-well assay format (mean ± s.e.m.; n=3). IC_50_ values and 95% CI are indicated. **(G)** Effect of the gametocytocidal hit compounds on asexual parasite proliferation using the 96-well assay format (mean ± s.e.m.; n=3). IC_50_ values and 95% CI are indicated.

This primary screen identified 987 compounds that caused a > 30% reduction in luminescence intensity compared to untreated controls (Fig. 2C, Dataset S1). Of these, 884 molecules were available for 8-point dose response experiments (concentration range: 0.3 nM to 5 µM) on stage V gametocytes using the same high throughput assay setup as explained above, as well as on HepG2 cells to assess toxicity against human cells (Fig. 2C, Dataset S1).

This validation screen identified 205 hits with half-maximal inhibitory concentrations (IC_50_) < 2 µM against stage V gametocytes, and of the 156 molecules available for SYBR Green I-based dose response assays on asexual blood stage parasites, around 70% also inhibited parasite replication (Dataset S1). As expected from the diversity-oriented nature of the main compound library, hit clustering did not reveal prominently enriched molecule classes (Dataset S1). Focusing on a final hit list of the 85 most active molecules (gametocytocidal IC_50_ < 0.5 µM) with a favorable selectivity index (SI > 10), 29 compounds displayed IC_50_ values below 100 nM against both stage V gametocytes and asexual parasites (Dataset S1). Many of these highly dually active molecules are known antibiotics, including seven polyether ionophores such as boromycin and monensin that are known for their antimalarial and gametocytocidal activities (Adovelande and Schrével, 1996; D’Alessandro *et al*., 2015; De Carvalho *et al*., 2022; Brancucci *et al*., 2024) (Fig. 2D). Remarkably, however, this set also included eight translation inhibitors, such as the tRNA synthetase-targeting molecules halofuginone and borrelidin that have previously been shown to target different parasite life cycle stages (Jain *et al*., 2015; Herman *et al*., 2015; Novita *et al*., 2024; Ishiyama *et al*., 2011). In contrast to these dually active compounds, several hits showed lower or no activity against asexual parasites, including six molecules identified as highly active in the gametocyte viability assay (IC_50_ < 100 nM), indicating gametocyte-selective activity (Dataset S1).

### The dually active compounds CR-1-31-B, ONX-0914, brusatol and halofuginone have potent transmission-blocking activity

We selected six chemically tractable hit molecules for further analyses, four with potent dual activity [halofuginone (cpd 209), CR-1-31-B (cpd 643), brusatol (cpd 210) and ONX-0914 (cpd 151)] and two that appeared to specifically target gametocytes (cpd 390 and cpd 464) (Fig. 2E). Halofuginone inhibits the parasite’s cytoplasmic prolyl-tRNA synthetase. CR-1-31-B is a synthetic analog of rocaglates that inhibit eukaryotic translation initiation factor eIF4A (Shen and Pelletier, 2020). ONX-0914 is an inhibitor of the mammalian immunoproteasome subunits β1i (low– molecular mass polypeptide 2, LMP2) and β5i (LMP7) (Muchamuel *u. a.*, 2009; Jenkins *u. a.*, 2021). Brusatol is known for inhibiting the NRF2 stress response pathway in human cancer cells (Xi *et al*., 2024). Activity against asexual parasites has previously been demonstrated for all four dually active molecules (Langlais *et al*., 2018; Cox *et al*., 2024; Van Truong *et al*., 2025), but their activity against the quiescent stage V gametocytes and their transmission to the mosquito vector has so far not been investigated in any great detail.

First, we performed dose response assays with independent compound batches on both parasite stages using the 96-well plate assay format to confirm the results obtained in the high-throughput validation screen (Figs. 2F-G, Dataset S1). All six molecules showed high activity against stage V gametocytes. Notably, however, CR-1-31-B only led to a partial reduction of the luminescence signal emitted from stage V gametocytes, leveling off approx. 30% below the signal obtained for the untreated controls (as also observed in the validation screen), indicating potential sex-specific activity (Fig. 2F). CR-1-31-B, halofuginone, brusatol and ONX-0914 also inhibited asexual proliferation in the single-digit nanomolar range, while cpd 390 and cpd 464 showed no activity (Fig. 2G).

To test whether the gametocytocidal effects observed in vitro translate into transmission-blocking activity, we performed standard membrane feeding assays (SMFAs). To this end, stage V gametocytes of the NF54-HGL strain that express a GFP-firefly luciferase fusion reporter (Vos *et al*., 2015) were exposed to the compounds for 48 hours at two concentrations (IC_50_ and 10x IC_50_) in duplicates. Untreated gametocytes and gametocytes treated with 10 µM dihydroartemisinin (DHA) were used as negative and positive controls for transmission blockade, respectively. Treated gametocytes were subsequently fed in duplicate to female *Anopheles stephensi* mosquitoes and eight days later the proportion of infected mosquitoes (infection prevalence) and the luminescence signals emitted from individual mosquitoes as a surrogate for the number of oocysts per mosquito (infection intensity) were quantified (Vos *et al*., 2015; Dechering *et al*., 2017). Strikingly, halofuginone, CR-1-31-B, brusatol and ONX-0914 were highly effective and prevented parasite transmission to mosquitoes already at the IC_50_ concentrations determined for stage V gametocyte killing (Fig. 3A). In contrast, cpd 390 and cpd 464 were inactive at both concentrations tested, suggesting these compounds were false positive hits in the HTS assay and inhibited luminescence emission rather than killing gametocytes. While the structures of both of these compounds do not show obvious similarities with luciferase inhibitors, their aromatic nature suggests they may absorb light emission through an inner filter effect (Simeonov *et al*., 2008; Thorne, Inglese and Auld, 2010). Functional gametocyte activation assays revealed that halofuginone, brusatol and ONX-0914 blocked the formation of both female and male gametes (Fig. 3B). In contrast, CR-1-31-B exclusively inhibited male gamete formation (Fig. 3B), which is consistent with the partial inhibition of luciferase reporter activity observed in the gametocytocidal assays (Fig. 2E). To confirm the sex-specific activity of CR-1-31-B, we probed the effect of its structural analog rocaglamide A (RocA), a natural compound of the rocaglate class (Rodrigo *et al*., 2012), on gamete formation. Similar to CR-1-31-B, RocA was inactive in the macrogamete formation assay but highly effective in preventing male gametocyte exflagellation (Fig. 3B).

Finally, we investigated whether CR-1-31-B, brusatol and ONX-0914 activity is compromised in parasites resistant to known antimalarial drugs. To this end, we tested *P. falciparum* strains NF54, RF12 (PH1263-C) (resistant to chloroquine, piperaquine, pyrimethamine, artemisinin) (Ross *et al*., 2018; Kanai *et al*., 2022) and Dd2 [resistant to chloroquine (Wellems *et al*., 1990), cycloguanil and pyrimethamine (Kreutzfeld *et al*., 2022)], as well as Dd2 lines engineered to carry point mutations conferring resistance to GNF156 (Magistrado *et al*., 2016), SJ557733 (Qiu *et al*., 2022), MMV019721, MMV084978 (Summers *et al*., 2022), MMV390048 (Paquet *et al*., 2017) and DSM265 (Mandt *et al*., 2019) in dose response assays on asexual parasites. All parasites showed similar sensitivity to CR-1-31-B, brusatol and ONX-0914 as NF54/iGP1_RE9H^pfs16^, demonstrating that the mechanisms of resistance for known antimalarial drugs do not confer cross-resistance to the hit compounds identified here (Table S1).

**Figure 3.**
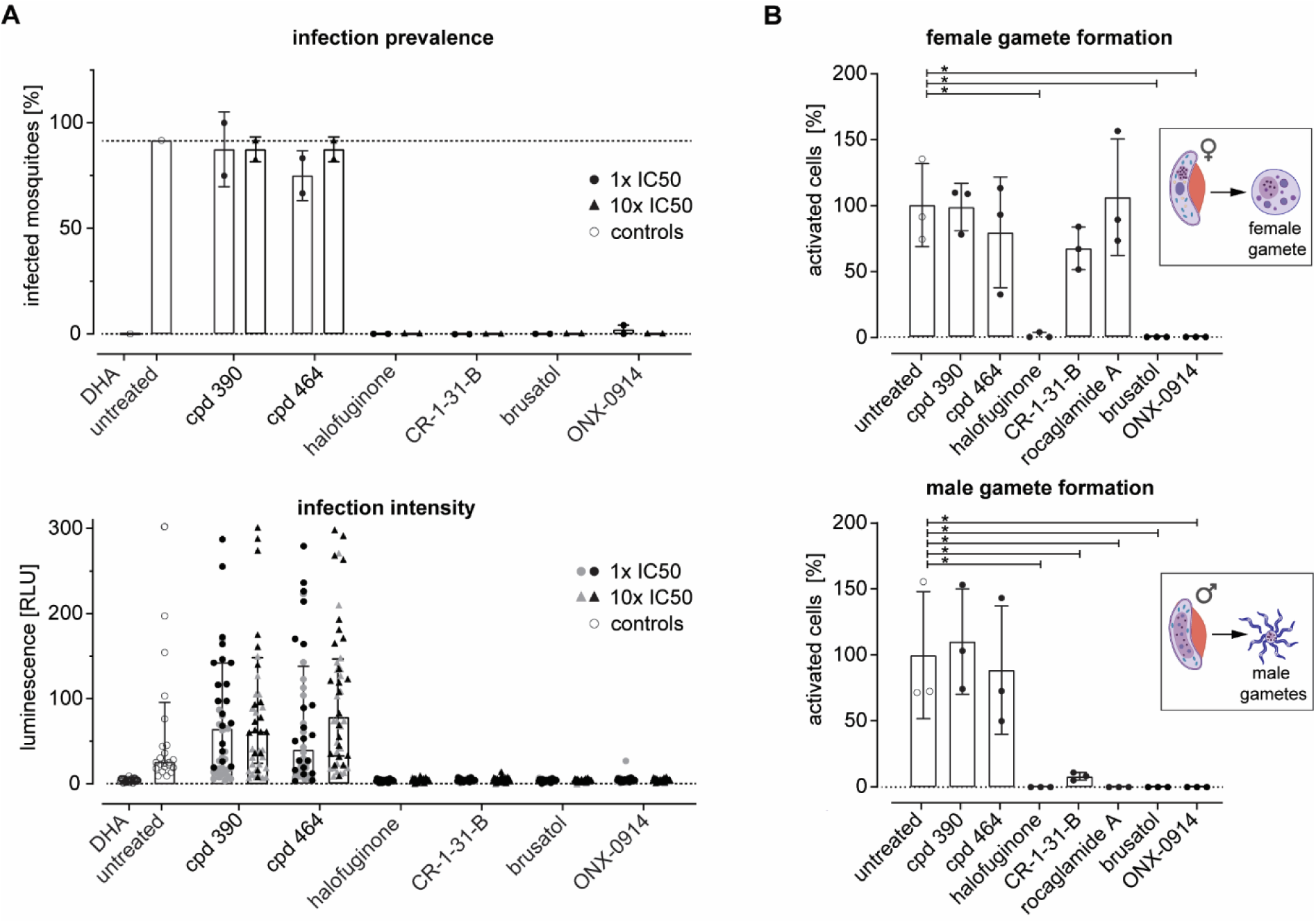
Halofuginone, CR-1-31-B, brusatol and ONX-0914 potently inhibit transmission to mosquitoes and female and/or male gametocyte activation. **(A)** Effect of selected hit compounds on parasite transmission to mosquitoes. The top panel shows the percentage of infected among all blood-fed mosquitoes (infection prevalence) (mean of duplicate feeds ± s.d.). Compounds were tested at their gametocytocidal IC_50_ and 10x IC_50_ concentrations using 24 mosquitoes per feed (12 mosquitoes were used for each feed on untreated control gametocytes). The bottom panel shows infection intensity (oocysts/mosquito), with each dot representing the oocyst load in individual mosquitoes as quantified by luminescence readout, and results from different feeds are presented in black and grey. Median and interquartile ranges are indicated; n=2. DHA, 10 µM dihydroartemisinin (positive control); untreated, 0.1% DMSO (negative control). **(B)** Effect of hit compounds on the formation of female gametes (top panel) and male gametes (bottom panel). Compounds were tested at 10x of their gametocytocidal IC_50_ concentrations. Data indicates the proportion of gametocytes forming gametes normalized to the mean of the untreated controls (mean ± s.d.; n=3). Asterisks mark statistically significant differences (unpaired, two-tailed t-test; p-value < 0.05).

### In vitro selection of parasites resistant to CR-1-31-B, brusatol and ONX-0914

To identify the molecular targets of CR-1-31-B, brusatol and ONX-0914, we selected asexual Dd2-B2 parasites (Eastman *et al*., 2011; Ross *et al*., 2018) for resistance and evaluated the effects of resistance mutations on gametocytocidal drug activities.

For each compound, we initiated resistance selection in triplicate experiments using 5-10x10^8^ asexual parasites and three to six times the IC_50_ determined for inhibition of asexual NF54 parasite replication (Fig. 2G). Two independent parasite populations each resistant to either CR-1-31-B (populations A and B) or brusatol (populations C and D), as well as one ONX-0914-resistant population (population E) were successfully retrieved between three to seven weeks under constant compound pressure. From each resistant population, we isolated parasite clones by limiting dilution and performed dose-response assays on asexual parasites (Fig. 4A, Table S2). Compared to the Dd2-B2 parent, CR-1-31-B-resistant clones derived from population A tolerated up to 2.8-fold higher compound levels (2.1-2.8-fold, three clones) and those derived from population B were completely insensitive to CR-1-31-B (three clones) (Fig. 4B). Similarly, brusatol-selected clones showed up to 19-fold (15.1-19-fold, four clones derived from population C) or over 100-fold decreased sensitivity to brusatol (five clones derived from population D) (Fig. 4C). Finally, all seven parasite clones originating from the ONX-0914-selected population (population E) showed up to 6.7-fold increased drug tolerance (5-6.7-fold) (Fig. 4D).

Next, we tested whether stage V gametocytes produced from the resistance-selected parasites were also less sensitive to the corresponding compounds. We chose one representative clone each derived from population B (CR-1-31-B-resistant clone B1), populations C and D (brusatol-resistant clones C5 and D1) and population E (ONX-0914-resistant clone E1) and assessed compound activity using a high content imaging-based readout of mitochondrial activity and cellular shape of MitoTracker- and SYBR Green I-stained cells (Kuehnel *et al*., 2023; Brancucci *et al*., 2025). Indeed, gametocytes derived from these resistant parental populations showed decreased sensitivity to the respective drugs by >100-fold (CR-1-31-B-resistant clone B1), 21.9-fold and >100-fold (brusatol-resistant clones C3 and D1, respectively) and 5.5-fold (ONX-0914-resistant clone E1), indicating shared targets and/or resistance mechanisms in asexual parasites and gametocytes (Fig. 4E-G).

**Figure 4.**
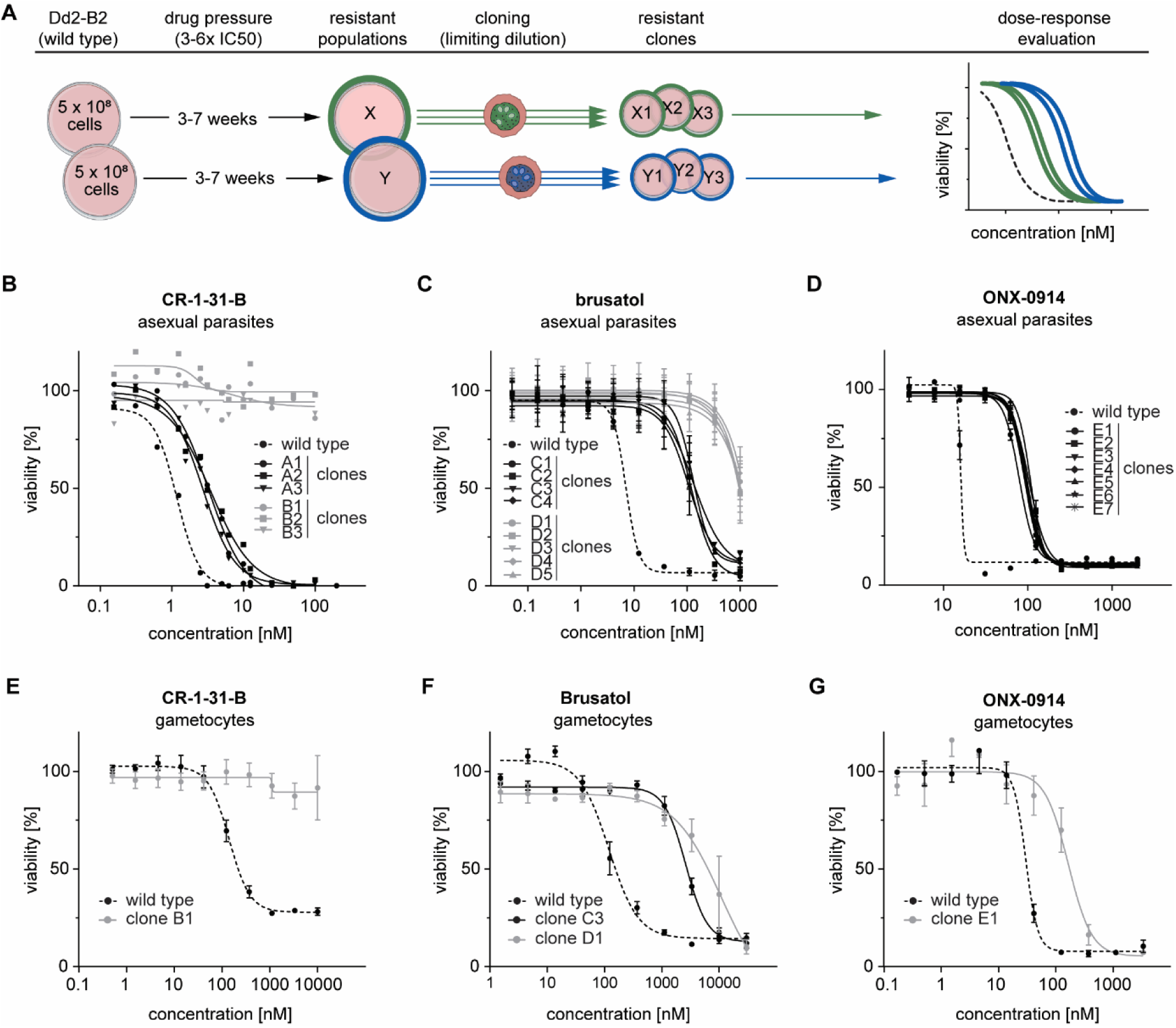
In vitro selection yields drug-resistant parasite lines. **(A)** Schematic representation of the resistance selection procedure. **(B-D)** Asexual parasites of clonal lines derived from resistance-selected Dd2-B2 populations show decreased susceptibility to CR-1-31-B (n=2) (B), brusatol (n=3) **(C)** and ONX-0914 (n=3) **(D)**. Data points represent the mean ± s.e.m., where applicable. **(E-G)** Stage V gametocytes of CR-1-31-B-resistant clone B1 **(E)**, brusatol-resistant clones C3 and D1 **(F)** and ONX-0914-resistant clone E1 **(G)** show decreased susceptibility to the corresponding compounds (mean ± s.e.m.; n=3).

### Whole genome sequencing identifies the putative targets of CR-1-31-B, ONX-0914 and brusatol

To investigate whether resistance to CR-1-31-B, brusatol and ONX-0914 was linked to mutations in genes encoding potential targets, we performed whole genome sequencing (WGS) of three clones each derived from parasite populations A and B (CR-1-31-B-resistant), C and D (brusatol-resistant) and E (ONX-0914-resistant). We also sequenced genomic DNA of the parental Dd2-B2 line harvested at both the start and the end of the selection period to allow excluding mutations that may have accumulated independent of the selection process. After identifying and annotating variants present in all sequenced clones, we filtered for high-confidence mutations present in all three resistant parasite clones sequenced per population, and excluded mutations also detected in the two Dd2-B2 wild-type genomes. In each case, only one missense SNP located in a protein-coding region passed these filters (Table S3).

All six clones derived from two independent CR-1-31-B-resistant populations carried a non-synonymous SNP in either exon 1 or 2 of the gene encoding *P. falciparum* PfeIF4A (PF3D7_1468700) (Fig. 5 and Table S3). The corresponding amino acid changes were associated with different levels of resistance: clones obtained from population A (2-3-fold increased tolerance) all featured a substitution of threonine at position 100 with alanine (Thr100Ala), while clones derived from population B (>100-fold resistance) carried a lysine residue instead of glutamine at position 185 (Gln185Lys). These findings are consistent with the fact that rocaglates and synthetic analogs including CR-1-31-B target eIF4A in humans (Chu and Pelletier, 2015) and have been shown to interact with recombinant PfeIF4A in vitro (Langlais *et al*., 2018).

All three population E-derived ONX-0914-resistant clones (5-7-fold increased tolerance) shared a SNP in the gene encoding the *P. falciparum* 20S proteasome β5 subunit (Pf20S β5, PF3D7_1011400), resulting in a substitution of methionine with isoleucine at position 105 (Met105Ile) (Fig. 5), which aligns well with the known activity of ONX-0914 against the human immunoproteasome β5i/LMP7 subunit (Muchamuel *u. a.*, 2009; Jenkins *u. a.*, 2021).

All three brusatol-resistant clones derived from the highly resistant population D (>100-fold) shared a unique SNP that results in an exchange of glutamic acid with aspartic acid at position 864 (Glu864Asp) in the putative large subunit rRNA methyltransferase (PF3D7_1354300) (Fig. 5). rRNA methyltransferases play a key role in ribosome biogenesis by methylating ribosomal RNA. Mutation or acquisition by plasmid transfer of these enzymes in bacteria has been reported to confer resistance to multiple inhibitors of protein synthesis, such as macrolides, phenicols, lincosamides and streptogramin A antibiotics (Madsen *et al*., 2005; Long *et al*., 2006; Pakula, Hansen and Vester, 2017). While brusatol is known for inhibiting the NRF2 stress response pathway in human cancer cells (Xi *et al*., 2024), this effect was proposed to be a downstream product of reduced ribosomal activity and thus decreased expression levels of NRF2 and other short-lived proteins (Harder *et al*., 2017). Indeed, a co-crystallization study performed with archaea-derived ribosomes revealed that the brusatol-analog bruceantin binds to and potentially blocks the peptidyl-transferase center (PTC), the catalytic core of the ribosome (Gürel *et al*., 2009). We thus considered that the mutation in the putative large rRNA methyltransferase PF3D7_1354300 identified in the highly brusatol-resistant D clones points toward a resistance mechanism involving the ribosome as a drug target. By contrast, we could not identify any consistent SNPs in the three clones derived from population C that showed moderate brusatol resistance (15-19-fold). We also found no indication for relevant gene copy number variations (CNVs) in these genomes (Fig. S2, Dataset S2), indicating that transcriptional and/or post-transcriptional changes may be responsible for increased brusatol tolerance in these clones.

In summary, our in vitro resistance selection experiments coupled with WGS successfully identified the likely molecular targets and mode-of-actions of the potent transmission-blocking compounds CR-1-31-B, brusatol and ONX-0914.

**Figure 5.**
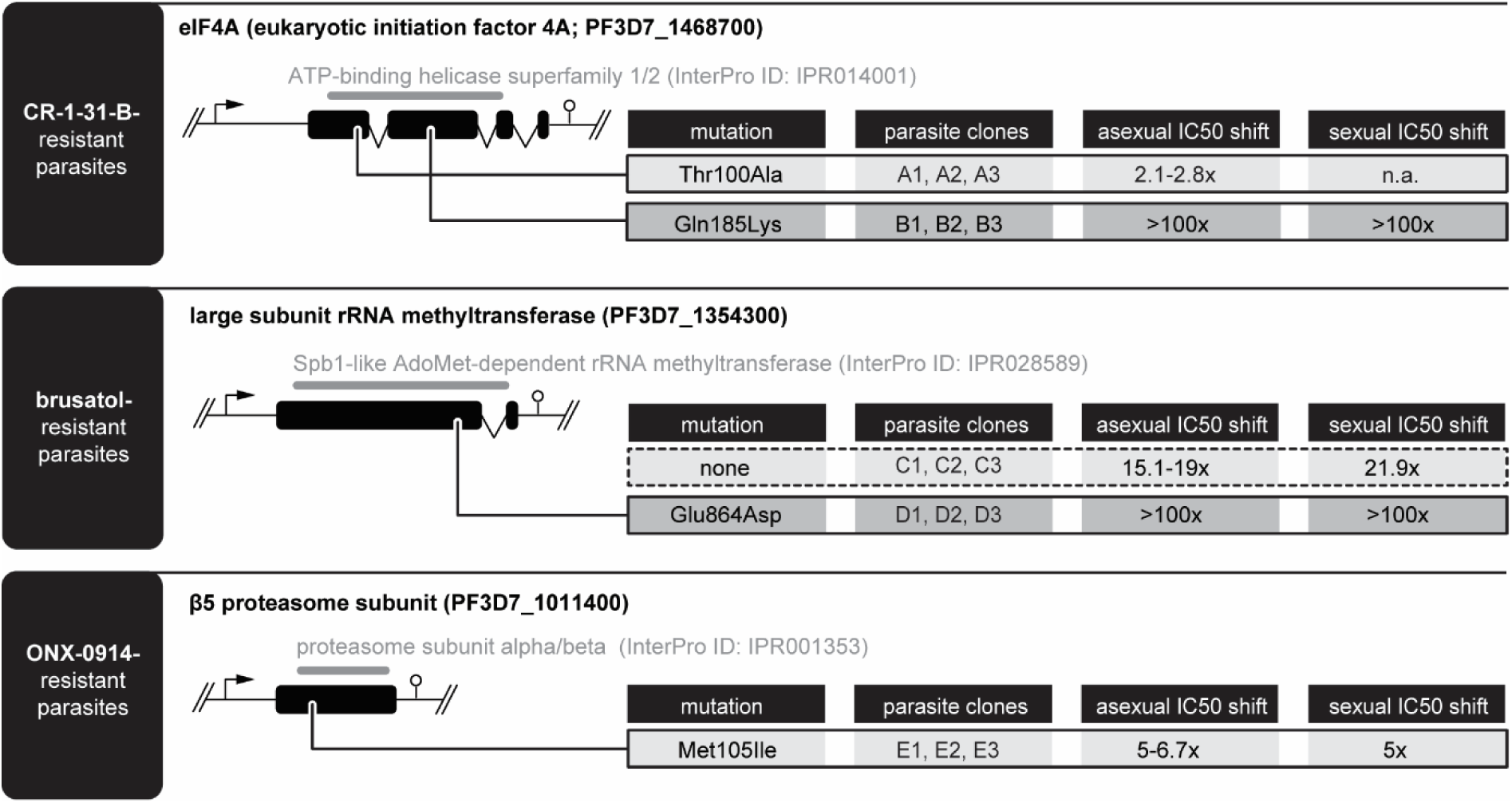
Clonal lines derived from resistance-selected parasites carry unique SNPs in putative target genes. Schematic maps showing the non-synonymous SNPs found in clonal Dd2-B2 lines selected in vitro for resistance to CR-1-31-B (upper panel), brusatol (center panel) and ONX-0914 (lower panel). Amino acid changes and drug sensitivity shifts in asexual and sexual parasites relative to the Dd2-B2 parent are indicated. Gene IDs, product descriptions and exon-intron structures are shown (www.plasmodb.org). InterPro domain annotations are shown in grey.

### Mutations in PfeIF4A and the Pf20S β5 proteasome subunit alter inhibitor binding affinity

We reasoned that parasite resistance to CR-1-31-B and ONX-0914 is likely explained by the single amino acid substitutions identified in their putative target proteins PfeIF4A and Pf20S β5, respectively. Using a combination of homology modeling and constrained docking, we therefore generated structure models of wild-type PfeIF4A and the Thr100Ala and Gln185Lys mutants in complex with CR-1-31-B. Similarly, we modeled wild-type Pf20S β5 and the Met105Ile mutant in complex with ONX-0914 in its pre-reactive, i.e. non-covalently bound state.

The model of PfeIF4A (Q8IKF0_PLAF7) in complex with CR-1-31-B was generated using the X-ray structure of human eIF4A-I (PDB ID: 5zc9, P60842) co-crystallized with silvestrol, an analog of RocA and CR-1-31-B (Fig. 6A). The PfeIF4A/CR-1-31-B model revealed that key interactions are preserved relative to the silvestrol-bound crystal structure of human eIF4A-I. Notably, residue Gln185 plays a central role in a hydrogen bond network as it interacts with the methoxyacetone group of CR-1-31-B (corresponding to the 1-(methylamino)propan-2-one group in silvestrol) on one side, and with the adenosine nucleotide and the RNA sugar backbone on the other side (Fig. 6A). Funnel metadynamics simulations indicated a significant difference in protein-ligand interactions strength between the wild-type PfeIF4A and the Gln185Lys mutant (Fig. S3). For wild-type PfeIF4A, the estimated binding free energy is –10.9 kcal/mol (Kd ≈ 11.9 nM), whereas in the mutant it is –4 kcal/mol (Kd ≈ 1.3 mM). These results strongly suggest that disruption of the Gln185-mediated hydrogen bond network is responsible for the >100-fold decrease in CR-1-31-B sensitivity observed experimentally. Like the Gln185 residue, Thr100 is located near the RNA-binding site and engages in a hydrogen bond network (Fig. 6A). This residue interacts with the backbone nitrogen of Leu103 and likely forms an additional water-mediated interaction with Leu321. However, unbiased molecular dynamic (MD) simulations did not reveal significant structural differences between the wild-type and Thr100Ala mutant target-inhibitor complexes, suggesting that any structural or functional impact of the mutation may be allosteric in nature and occur on a longer timescale that remains undetected by standard MD simulations. This absence of pronounced effects at the structural level aligns with the markedly lower level of CR-1-31-B resistance observed for the Thr100Ala (>2-fold) compared to the Gln185Lys mutants (>100-fold).

**Figure 6.**
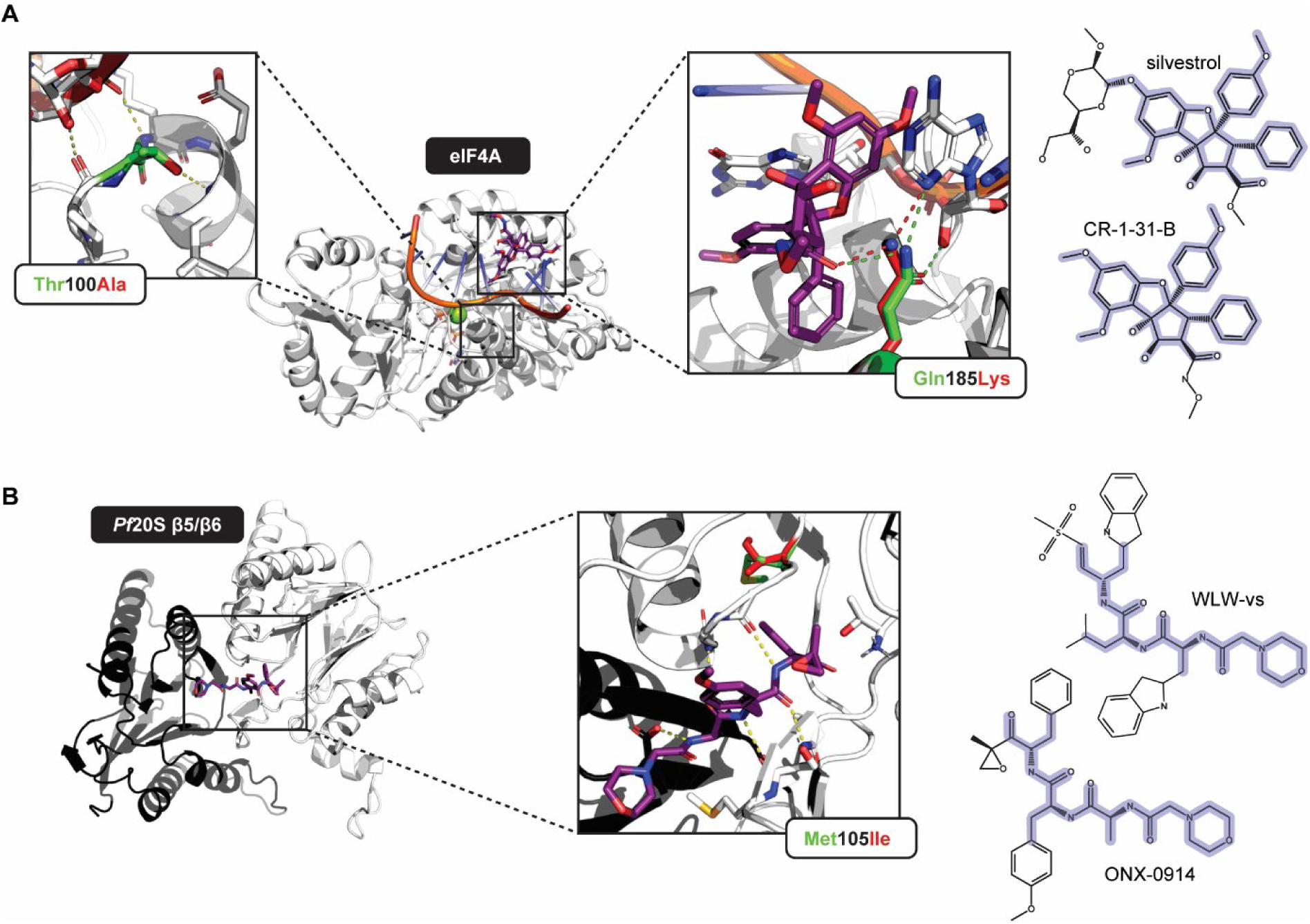
Interaction modelling reveals the targets of CR-1-31-B and ONX-0914. **(A)** The Gln185Lys mutation in PfeIF4A, and to a lesser extent also the Thr100Ala mutation, reduce binding affinity for CR-1-31-B. The compound is shown together with its structural analog silvestrol, with overlaps highlighted in blue. **(B)**. The Met105Ile mutation in the Pf20S β5 subunit alleviates interactions with ONX-0914. Hydrogen bonds (dashed lines) between the compound (purple) and amino acid residues in their wild-type (green) and mutated configuration (red) are shown. The compound is shown together with its structural analog WLW-vs, with overlaps highlighted in blue.

To study the interaction of ONX-0914 with Pf20S β5 subunit, we made use of the cryo-EM structure of the *P. falciparum* 20S proteasome complexed with inhibitor WLW-vs (Hsu *et al*., 2023), with the ligand replaced by ONX-0914 (Hsu *et al*., 2023) (Fig. 6B). Similar to related epoxyketone compounds showing activity against *P. falciparum* asexual parasites (e.g. J-78 or J-80) (Almaliti *et al*., 2023; Deni *et al*., 2023), ONX-0914 is a peptide-based inhibitor that features a reactive epoxide warhead at the N-terminus and a morpholino group at the C-terminus (Muchamuel *et al*., 2009). Based on the crystal structure of ONX-0914 complexed with the mouse immunoproteasome (de Bruin *et al*., 2014), the epoxide group likely covalently binds the catalytic Thr1 residue in the Pf20S β5 subunit. Relative free energy calculations of ONX-0914 binding performed with Desmond and BioSimSpace predicted a reduction in binding affinity of 0.9 kcal/mol and 2.0 kcal/mol, respectively, for the Met105Ile mutant compared to wild-type Pf20S β5. Analysis of the MD trajectories for the two end states of the relative free energy calculations, i.e. for the wild-type and Met105Ile systems, revealed weakened stabilizing interactions involving the C-terminal region of the extended loop–β-hairpin–loop motif (residues 77-95) as well as residues Ala109 and Leu113. This loss of stabilization leads to increased flexibility of residues 77-95 and 105-110 (Fig. S4). As a consequence, ligand interactions with the backbone and side chain of Ser81 are weakened, with the hydrogen-bond occupancy decreasing from 1.65 in wild-type Pf20S β5 to 1.0 in the Met105Ile mutant. In addition, hydrogen-bond occupancies involving Gly107, Ala110, and Ala111 are reduced by 12%, 30%, and 34%, respectively. Most notably, the hydrogen-bonding frequency to the catalytic Thr1 is diminished by 76%, substantially lowering the probability of covalent reaction with the ONX-0914 warhead and providing a mechanistic explanation for the experimentally observed reduction in binding affinity.

### An atypical rRNA base methylation in the ribosomal A-site is linked to brusatol resistance

Parasites carrying a Glu864Asp mutation in the putative large subunit rRNA methyltransferase PF3D7_1354300 are highly insensitive to brusatol (>100-fold resistance compared to parental Dd2-B2 parasites). A template search revealed that PF3D7_1354300 (accession number XP_001350255) is closely related to the 27S pre-rRNA (guanosine(2922)-2’-O)-methyltransferase SPB1 of *Saccharomyces cerevisiae* (accession number P25582). PF3D7_1354300 shares 39.11% sequence identity (E-value of 2e-72) with ScSPB1 and 38.73% (E-value of 2e-81) with the human homolog pre-rRNA 2’-O-ribose RNA methyltransferase FTSJ3 (accession number Q8IY81), and the Glu864Asp mutation is located in the conserved C-terminal region (Fig. S5). Both ScSPB1 and human FTSJ3 play important roles during ribosome biogenesis but are absent from mature ribosomes (Kater *et al*., 2017; Vanden Broeck and Klinge, 2023). Studies in yeast have shown that ScSPB1 is an essential structural component of nucleolar pre-60S intermediates and responsible for methylating the universally conserved nucleotide G2922 in the 25S rRNA (corresponding to G3281 in *P. falciparum* 28S rRNA) located in the catalytic PTC adjacent to the aminoacyl tRNA-acceptor site (A-site) (Figs. S5, S6 and Table S4) (Lapeyre and Purushothaman, 2004; Taoka *et al*., 2018). The ribosomal A-site is overall highly conserved in all kingdoms of life (Tirumalai *et al*., 2021) and known to be targeted by bruceantin in archaeal organisms (Guru *et al*., 1983; Gürel *et al*., 2009). Bruceantin differs from brusatol only by two methyl groups at the ester side chain. To the best of our knowledge, neither of these compounds was ever proposed to interact with methyltransferases directly. In light of these findings, the *P. falciparum* putative large subunit rRNA methyltransferase PF3D7_1354300 (henceforth termed putative PfRLM) is unlikely the direct target of brusatol. Instead, we speculated that brusatol resistance may be linked to differential 28S rRNA base modifications within the PTC, as a result of altered enzymatic activity of the putative PfRLM Glu864Asp mutant.

To investigate whether the 28S rRNA of brusatol-resistant parasites is indeed differentially methylated, we used an Oxford Nanopore Technologies (ONT)-based approach capable of detecting post-transcriptional modifications within native rRNA molecules (Smith *et al*., 2019; Delgado-Tejedor *et al*., 2024). The nanopore current generated by base G3281 differed markedly between 28S rRNA isolated from parasites and in vitro-transcribed (IVT) control 28S rRNA, demonstrating that this nucleotide is modified in native 28S rRNA in *P. falciparum*, similar to the ScSPB1-targeted base equivalent G2922 in yeast 25S rRNA (Figs. S5, S6, Table S4). While we could not observe differential modification of G3281 between brusatol-sensitive and -resistant parasites, several 28S rRNA bases positioned in the vicinity of the A-site showed differences in normalized ONT-sequencing current in brusatol-resistant parasites (Fig. 7A). Specifically, bases A2665-C2667 in helix 72 of the large subunit domain V were prominently affected in clone D1 parasites that express the mutated form of putative PfRLM and display >100-fold resistance to brusatol (Fig. 7A) (for positions in *S. cerevisiae, E. coli, A. thaliana* and *H. sapiens*, see Fig. S6 and Table S4). Intriguingly, clone C1 parasites, which express the wild-type putative PfRLM and show a less pronounced decrease in brusatol-sensitivity (15-19-fold), carry the same atypical 28S rRNA modifications, although at lower frequencies compared to the highly resistant clone D1 (Fig. 7A). These results imply that brusatol-resistant parasites catalyze the non-canonical modification of 28S rRNA bases in the large ribosomal subunit. If the putative PfRLM is indeed a functional homolog of ScSPB1, the modifications at the 28S rRNA are expected to represent 2’-O-ribose methylations. Because Nanopore sequencing cannot discriminate between different types of post-transcriptional base modifications, we used Nm-VAQ, an approach capable of identifying methylated RNA bases (Tang *et al*., 2024). Nm-VAQ exploits the fact that only 2’-O-ribose methylated RNA is protected from RNase H-mediated cleavage when hybridized to complementary DNA (DNA/RNA chimera) (Fig. 7B). Nm-VAQ revealed that base C2667 of the 28S rRNA was indeed protected from RNase H activity in drug-resistant parasites, demonstrating that this base carries a novel 2’-O-ribose methylation (Figs. 7B and S7). By contrast, we did not find convincing evidence for methylation of bases A2665-2666, indicating that their ONT-signature was influenced by the modification of their neighbor at position 2667 (Leger *et al*., 2021). Together, these findings indicate that the Glu864Asp mutant of the putative PfRLM catalyzes the unusual 2’-O-ribose methylation of C2667 in 28S rRNA helix 72 in brusatol-resistant parasites.

In Archea, bruceantin binds to the PTC close to the A-site tRNA acceptor arm, which perturbs the geometry of the PTC, hindering elongation by sterically blocking peptidyl transfer (Gürel *et al*., 2009). While the atypical methylation we identified in helix 72 of the *P. falciparum* 28S rRNA does not directly interface with the binding site for brusatol, helix 72 forms contacts with helix 89, including A3179 and C3180 in the PTC, both of which are important for the interaction with bruceantin (A2820 and C2821 in *S. cerevisiae*) (Gürel *et al*., 2009) (Fig. 7C). In this context, it is worth noting that we also identified potent gametocytocidal activity for a second A-site-targeting compound, homoharringtonine (see Fig. 2D; IC_50_ = 3.8 nM; 95% CI: 0.01-10.1 nM). Despite being structurally unrelated to brusatol, homoharringtonine interacts with the same A-site-defining rRNA base in the archaea *H. marismortui* (U2871 in *S. cerevisiae*) (Gürel *et al*., 2009). We therefore tested whether brusatol-resistant parasites are also insensitive to homoharringtonine but could not detect cross-resistance (Fig. 7D), indicating that the brusatol resistance mechanism does not interfere with the binding affinity of homoharringtonine. Combined with the fact that homoharringtonine requires less space proximal to A2820 and C2821 (coordinates from *S. cerevisiae* 25S rRNA), these results further support the hypothesis that direct interactions between helix 89 and the atypically methylated helix 72 are causative for brusatol-resistance in *P. falciparum*.

**Figure 7.**
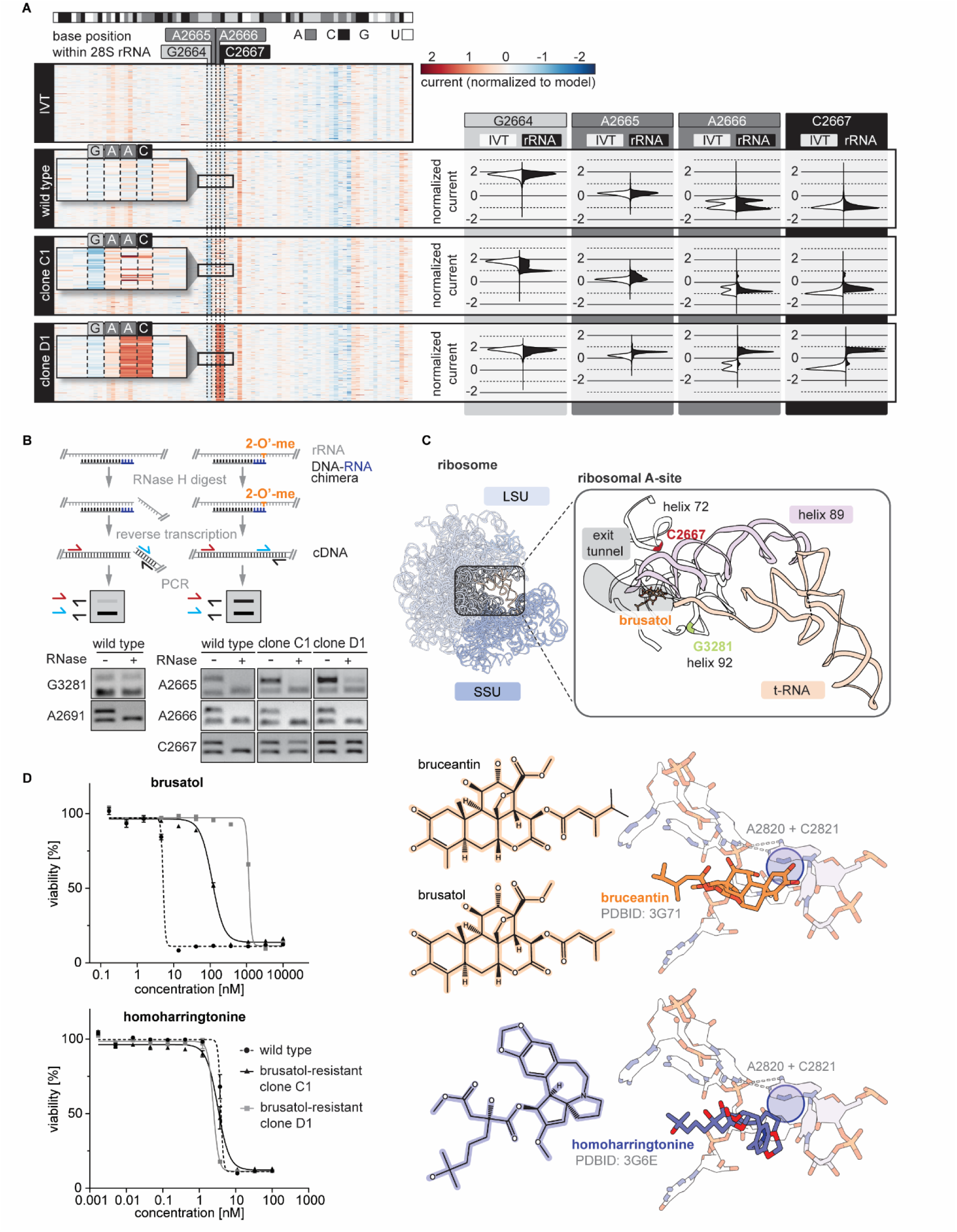
Atypical 28S rRNA methylations at the ribosomal A-site confer resistance against brusatol. **(A)** Four atypical 28S rRNA modifications are detected in moderately (clone C1) and highly (clone D1) brusatol-resistant parasites. The heatmap shows the normalized current measured by Nanopore sequencing of *in vitro*-transcribed (IVT) and native 28S rRNA isolated from brusatol-sensitive (wild-type) and -resistant parasites (clones C1 and D1). Reads were aligned using Uncalled4 (Kovaka *et al*., 2025). The color gradient represents the difference between the detected and expected normalized currents. rRNA base positions are indicated. Violin plots show the distribution of normalized current measured from IVT (white) and native (black) 28S rRNA molecules. **(B)** Nm-VAQ reveals novel 2’-O-methylation sites at position C2667 in brusatol-resistant parasite clones C1 and D1. 2’-O-methylated RNA protects RNA/DNA substrates from RNaseH-digestion and, following reverse transcription, can be PCR-amplified using primers indicated in red and blue. Amplification of a shorter PCR fragment that does not include the base of interest (using the primer highlighted in blue) served as a control. **(C)** Upper panel: Schematic of the small (18S, SSU) and large (28S + 5.8S, LSU) rRNAs from *P. falciparum* (PDBID: 9BUT) superimposed with *E. coli* tRNA (PDBID: 7K00) (beige). The zoom-in shows the arrangement of rRNA helices that surround the A-site portion of the PTC superimposed with brusatol at the start of the ribosomal exit tunnel. Methylated G3281 in helix 92 (highlighted in green) and the atypically methylated base C2667 in helix 72 of brusatol-resistant parasites (highlighted in red) are indicated. Lower panel: Chemical structures of bruceantin and brusatol (which differ by only two methyl groups at the ester side chain) (top) and homoharringtonine (bottom) and their binding sites at the ribosomal A-site (highlighted by blue circles) are shown in the context of the intercalating 25S rRNA bases A2820 and C2821 of *S. cerevisiae*. **(D)** Brusatol-resistant parasites show no cross-resistance to the A-site targeting drug homoharringtonine (mean ± s.e.m.; n=3).

## DISCUSSION

Here, we engineered the NF54/iGP1_RE9H^pfs16^ parasite line suitable for the mass production of synchronous *P. falciparum* gametocytes expressing the RE9H firefly luciferase under the control of the pfs16 promoter. We used NF54/iGP1_RE9H^pfs16^ stage V gametocytes to probe a large diversity-oriented compound library under high throughput conditions and identified several dozen compounds targeting quiescent stage V gametocytes with IC_50_ values in the low nanomolar range. For three of the most effective and chemically tractable compounds (CR-1-31-B, ONX-0914, brusatol), we (i) demonstrated their potent dual activity against stage V gametocytes and asexual parasites; (ii) investigated their effect on male and female gamete formation; (iii) confirmed their transmission-blocking activity in mosquito feeding assays; (iv) identified their cellular targets using resistance selection coupled to whole genome sequencing; and (v) modeled their interactions with their respective targets. Our efforts toward target deconvolution of the highly active compounds CR-1-31-B, ONX-0914 and brusatol revealed that stage V gametocytes share with asexual blood stages a particular vulnerability to proteostasis-targeting molecules.

Among the many different gametocytocidal drug assay formats applied to date, only a few are compatible with the screening of large compound libraries under high-throughput conditions (Plouffe, 2016; Sun *et al*., 2017; Delves *et al*., 2018; Paonessa *et al*., 2022). In their recent HTS campaign of >120,000 molecules, Paonessa and colleagues also used non-purified luciferase-expressing gametocytes in conjunction with the non-lysing D-luciferin substrate, which greatly simplified the production of suitable late stage gametocyte preparations and cellular viability readout (Paonessa *et al*., 2022). While our HTS assay developed here employs a comparable experimental setup, the high sexual conversion rates achieved by NF54/iGP1_RE9H^pfs16^ parasites substantially improve both gametocyte yield and synchronicity. As a result, we were able to screen >50,000 compounds against pure mature stage V gametocytes on a single day, requiring only 60 ml of gametocyte culture. The compounds screened in this study derived from different non-proprietary Novartis libraries and included a collection of almost 49,000 diversity-oriented compounds, 841 compounds with known modes of action, as well as 2475 natural products. Of the 884 primary hits evaluated in dose-response assays, 85 displayed IC_50_ values below 0.5 μM and selectivity indices above 10. Forty-four of these compounds also target asexual parasites at submicromolar IC_50_ concentrations and 29 of these dually active molecules killed both parasite stages at IC_50_’s below 100 nM. In contrast, several primary hits showed high activity in the gametocyte viability assay but were inactive against asexual parasites. However, gamete formation and SMFA experiments performed with two of these molecules identified them as false-positive gametocytocidal hits. We consider inhibitory effects on luciferase activity the most likely explanation for this outcome, but a more systematic analysis of this group of compounds is required to figure out whether any of these molecules truly possess gametocyte-specific activity.

eIF4A is a highly conserved translation initiation factor and part of the mRNA cap-binding complex eIF4F that regulates recruitment and 5’ UTR scanning of the 43S ribosomal pre-initiation complex (PIC). As an ATP-dependent DEAD-box RNA helicase, eIF4A resolves RNA secondary structures and RNA-protein interactions to facilitate efficient PIC scanning and translation initiation (Andreou and Klostermeier, 2013). Rocaglates, including CR-1-31-B, act by clamping eIF4A onto their target mRNAs, particularly at purine-rich stretches, thereby inhibiting translation of affected mRNAs (Iwasaki, Floor and Ingolia, 2016; Iwasaki *et al*., 2019; Shen and Pelletier, 2020; Shichino and Iwasaki, 2022). Although rocaglates are primarily known for their potent activity against many types of cancers, they have since also proven effective against pathogenic fungi, protozoan parasites (e.g. *Plasmodium* spp., *Cryptosporidium parvum*) and viral infections (e.g. SARS-CoV2) (Shichino and Iwasaki, 2022). Langlais and colleagues previously reported potent antimalarial activity of CR-1-31-B against asexual *P. falciparum* parasites and in vivo in *P. berghei*-infected mice (Langlais *et al*., 2018). Their study further demonstrated that CR-1-31-B increases the binding affinity of recombinant PfeIF4A with RNA and reduces translation efficiency in treated parasites (Langlais *et al*., 2018), similar to what was observed in model organisms (Chu *et al*., 2016; Iwasaki, Floor and Ingolia, 2016; Shen and Pelletier, 2020; Shichino and Iwasaki, 2022). Here, we showed that CR-1-31-B is also active against stage V gametocytes and displays potent transmission-blocking activity in the low nanomolar range. Intriguingly, however, our combined gametocytocidal and gamete formation assays revealed that CR-1-31-B, as well as the natural analog RocA, are only active against mature male but not female gametocytes. The sex-selective effect of these molecules therefore suggests that male gametocytes are more dependent on PfeIF4A-dependent translation initiation compared to females. Alternatively, immediate effects on females may be masked by DOZI-mediated translational repression – a scenario in which maternal effects of CR-1-31-B treatment would only become apparent following zygote formation.

The solved crystal structure of a complex formed by human eIF4A·RocA·RNA·ATP-analog provided a detailed understanding of how rocaglates clamp RNA onto eIF4A (Iwasaki *et al*., 2019). RocA binds in a cavity at the eIF4A-RNA interface where it interacts non-covalently via aromatic stacking and hydrogen bond formation with two adjacent purines in the RNA molecule and Phe163 and Gln195 in the RNA-binding pocket of eIF4A (Iwasaki *et al*., 2019). Phe163 plays a pivotal role in the sensitivity of eIF4A to rocaglate-mediated functional inhibition. In species naturally resistant to rocaglates, such as plants of the genus *Aglaia* that produce these secondary metabolites or *Ophiocordyceps sp. BRM1*, a pathogenic fungus of *Aglaia* spp., Phe163 is substituted with a non-aromatic Leu or Gly, respectively (Iwasaki *et al*., 2019; Chen *et al*., 2023; Obermann *et al*., 2023). Indeed, a systematic phylogenetic analysis of over 350 eIF4A orthologs across all eukaryotes discovered over a dozen naturally occurring substitutions at the position corresponding to Phe163 in human eIF4A, and those linked to rocaglate tolerance exist in species of several major lineages (Obermann *et al*., 2023). Interestingly, CR-1-31-B resistance in the *P. falciparum* clones selected in our study was not conferred by a mutation of this site (Tyr153 in PfeIF4A) but rather a substitution of the residue equivalent to Gln195 in human eIF4A (Gln185Lys in PfeIF4A). Unlike Phe163, Gln195 is strictly conserved among eIF4A orthologs (Obermann *et al*., 2023) and to our knowledge, rocaglate resistance via mutation of this residue has previously only been reported for the yeast eIF4A ortholog TIF1 in mutants selected for growth in the presence of Roc-N (Sadlish *et al*., 2013).

In contrast to CR-1-31-B, brusatol blocked the formation of male as well as female gametocytes. In human cells, brusatol is best known for inhibiting the NRF2 (nuclear factor erythroid 2-related factor 2) oxidative stress response pathway present in higher eukaryotes (Olayanju *et al*., 2015). Noteworthy, Harder *et al* proposed that this activity results from a general repression of translation that disproportionately affects proteins with short half-lives, including NRF2, rather than targeting NRF2 directly (Harder *et al*., 2017). Brusatol was recently linked to gametocytocidal and transmission-blocking activity in the *P. berghei* malaria mouse model (Cox *et al*., 2024). In this study, a 30-minute exposure to 100 nM brusatol blocked exflagellation of male gametocytes, while genome replication and axoneme formation remained unaffected. Female gametocytes were responsive to brusatol as well: following a two-hour drug pulse, macrogametes failed to express Pfs25, one of the first proteins expressed after mRNA release from DOZI-mediated repression (Tomas, 2001; Mair *et al*., 2006), which led the authors to speculate about a defect in protein synthesis in brusatol-treated cells (Cox *et al*., 2024). Cox and colleagues also confirmed activity of brusatol against *P. falciparum* immature gametocytes and male gametocyte exflagellation. In line with the view that brusatol targets translation, our WGS analysis revealed that a single amino acid substitution in the putative large subunit rRNA methyltransferase PfRLM (PF3D7_1354300) – an enzyme likely involved in ribosome biogenesis – renders parasites highly insensitive to the drug. Interestingly, in the archaeon *Haloarcula marismortui*, the brusatol analog bruceantin was found to inhibit translation by competing with the amino acid side chains of transfer RNA (tRNA) at the ribosomal A-site (Gürel *et al*., 2009). In yeast, the A-site contains the 28S rRNA base G2922 that is methylated by ScSPB1, the homolog of the putative *P. falciparum* PfRLM enzyme. Using ONT sequencing, we confirmed the presence of a Pf28S rRNA base modification at G3281 (corresponds to G2922 in *S. cerevisiae* 28S rRNA) in both wild type Dd2-B2 and brusatol-resistant parasites carrying the Glu864Asp mutation in the putative PfRLM. Intriguingly however, brusatol-resistant parasites derived from the highly resistant population D (IC_50_ shift >100-fold) carried a prominent modification at C2667 in the Pf28S rRNA molecule that is absent in wild type parasites. Likewise, this rRNA base was also modified at a lower level in parasites that are only moderately resistant to brusatol and express wild-type PfRLM (population C; IC_50_ shift = 15-19-fold). The G3281 base target of PfRLM is positioned in helix 92 and thus relatively far away from G2665-G2667 in helix 72 in the mature ribosome (Vanden Broeck and Klinge, 2023). In human cells, FTSJ3 associates dynamically with the pre-ribosomal complex and helices 72 and 92 are predicted to fold simultaneously with FTSJ3 binding during maturation (Vanden Broeck and Klinge, 2023). Provided that PfRLM interacts with pre-ribosomal complexes in a similar manner, it is conceivable that the mutated enzyme deposits methyl groups at helix 72 and thereby dislocates helix 89 at the PTC (see Fig. 7C). Notably, *H. marismortui* 23S rRNA bases A2486 and C2487 (A3179 and C3180 in Pf28S) in helix 89 play important roles for PTC activity as well as for the binding of the brusatol analog bruceantin to archaeal ribosomes (Gürel *et al*., 2009). We thus propose that the mechanism of brusatol resistance involves altered base-specificity of the PfRLM mutant and the consequent methylation of C2667 in Pf28S rRNA helix 72, ultimately lowering the binding affinity of brusatol to the ribosome without affecting translation. The fact that C2667 is also modified in parasites moderately resistant to brusatol strongly supports this scenario. How the C2667 base becomes methylated in these parasites expressing wild type PfRLM remains to be determined.

Noteworthy, our screen also identified high gametocytocidal activity for the ribosome-targeting drug homoharringtonine. This compound also binds to the large ribosomal subunit of *H. marismortui* in a manner similar to that of bruceantin (Gürel *et al*., 2009). Specifically, X-ray crystallography revealed that both homoharringtonine and bruceantin interact with the same A-site-defining 23S rRNA bases (A2486, C2487 in *H. marismortui*) and almost completely fill the A-site cleft, thereby preventing access of incoming aminoacyl-tRNAs (Gürel *et al*., 2009). Besides being polycyclic, bruceantin and brusatol are structurally unrelated to homoharringtonine and we indeed observed that the atypical Pf28S rRNA base modifications observed in brusatol-resistant parasites do not compromise the activity of homoharringtonine. The differences of these compounds regarding their structure and target interaction may offer opportunities for future structure-activity relationship (SAR) studies of ribosome-targeting drugs.

Alterations of ribosome-targeting binding sites, specifically through rRNA modifications, are well-studied in the context of antibiotic resistance and occur in both antibiotic-producing and pathogenic bacteria (Skinner, Cundliffe and Schmidt, 1983; Long *et al*., 2006; LaMarre, Howden and Mankin, 2011; Bhujbalrao and Anand, 2019; Osterman, Dontsova and Sergiev, 2020). In *E. coli* for instance, loss of the rRNA methyltransferase KsgA, which is responsible for the methylation of two neighboring adenosines in the 16S rRNA, confers resistance to the antibiotic kasugamycin (Van Buul and Van Knippenberg, 1985). The C8 carbon of the highly conserved 23S rRNA A2503 base in bacteria can be methylated by the methyltransferase Cfr, which confers resistance to eight classes of ribosome-targeting drugs, including tiamulin. Mutations in the amino acid sequence and promoter of Cfr increase the stability and activity of the enzyme, ultimately increasing methylation of A2503 and thereby causing antibiotic resistance (Tsai *et al*., 2022). Interestingly, the bacterial housekeeping rRNA methyltransferase RLmN also targets A2503 in the 23S rRNA, though in this case the methylation occurs at the C2 position and when methylation is lost due to a mutated RLmN, bacteria become more resistant (Stojković *et al*., 2016). To the best of our knowledge, altered specificity of rRNA methyltransferases was thus far not linked to drug resistance in *Plasmodium* spp. or in any other eukaryote. Considering the high conservation of ribosomal structure and maturation across all organisms (Wilson and Doudna Cate, 2012; Vanden Broeck and Klinge, 2024), however, ribosome-targeting molecules likely share a common mode of action in different organisms. We therefore believe the brusatol resistance mechanism we discovered here for *P. falciparum* provides valuable information about the drug’s mode of action and may thus also help clarify the anti-tumorigenic activity of brusatol and other A-site-targeting drug candidates in human cells (reviewed in Xi *et al*., 2024).

Proteasome-targeting compounds block the cell’s ability to degrade proteins via the ubiquitin-proteasome system, thereby triggering ER stress and the unfolded protein response, ultimately leading to cell death (Thibaudeau and Smith, 2019). Proteasome inhibitors have gained a lot of attention due to their activity against multiple myeloma, lymphoma and autoimmune diseases (Fricker, 2020; Wang *et al*., 2021), but are also well-known for their potent parasiticidal effects against different *Plasmodium* species (Rosenthal, 2023). In addition, proteasome-targeting drugs could be ideal candidates for antimalarial combination therapy with artemisinins, as the accumulation of damaged and unfolded proteins caused by artemisinin treatment cannot be dealt with when the proteasome is simultaneously inhibited (Dogovski *et al*., 2015; Stokes *et al*., 2019).

The *P. falciparum* core proteasome conforms to a typical 20S eukaryotic configuration (Wang *et al*., 2015), but was suggested to show a larger binding pocket in the chymotrypsin-like catalytic subunit β5, offering potential for developing parasite-specific inhibitors (Yasir *et al*., 2025). To date, several dozen covalent and non-covalent proteasome inhibitors targeting the β5 subunit have been demonstrated to kill asexual blood stage parasites with sub-micromolar to low-nanomolar potencies (Gantt *et al*., 1998; Stokes *et al*., 2019; Almaliti *et al*., 2023; Deni *et al*., 2023; Hsu *et al*., 2023; Rosenthal, 2023), a subset of which have also been tested against gametocytes showing promising activity, including the epoxyketones epoxomicin and carmaphycin B (Czesny *et al*., 2009; Aminake *et al*., 2011; Lelièvre *et al*., 2012; LaMonte *et al*., 2017; Kirkman *et al*., 2018; Xie *et al*., 2021; Van Truong *et al*., 2025). Here, we showed that ONX-0914, another epoxyketone also referred to as PR-957 with recently demonstrated activity against asexual stages (Van Truong *et al*., 2025), has high and nearly equipotent activity against asexual parasites and stage V gametocytes and effectively blocks gametocyte transmission at low nanomolar concentrations. We also showed via in vitro resistance selection, WGS and structural modeling that ONX-0914 targets the β5 subunit of the parasite 20S proteasome, consistent with the conserved mode of action of epoxyketone proteasome inhibitors (Wang *et al*., 2021). Notably, the Met105Ile mutation in the β5 subunit of ONX-0914-resistant parasites lowered their sensitivity only slightly (5-6.7-fold), such that dual activity in the nanomolar range is still retained in resistant parasites. Interestingly, the Met105 residue was also mutated in parasites selected for resistance to the epoxyketone J-80 (Met105Arg/Val) (Deni *et al*., 2023). J-80 is a synthetic analog of carmaphycin B with optimized pharmacological characteristics leading to a > 100-fold higher selectivity for the parasite compared to the human β5 subunit (LaMonte *et al*., 2017; Almaliti *et al*., 2023; Deni *et al*., 2023). On the contrary, ONX-0914 was designed to specifically target the β5i/LMP7 subunit of the human immunoproteasome and exerts more than 20-fold selectivity over the β5 subunit of the constitutive proteasome in leukemic human T-lymphoblasts (Muchamuel *et al*., 2009; Huber *et al*., 2012; Basler *et al*., 2015). These examples underscore that small structural changes allow refining engagement of this class of molecules with the β5 active site in different configurations, supporting a broader strategy of developing dual-active compounds specifically tailored to inhibiting the parasite β5 subunit, while avoiding cross-reactivity with human isoforms.

In conclusion, we developed an efficient and robust HTS assay specifically designed for the identification of compounds able to kill mature quiescent stage V gametocytes. With less than 1 µl of gametocyte culture required per test condition, this assay is highly suitable for the screening of comprehensive chemical libraries. Of the 51,185 compounds screened here, 29 validated hits showed highly potent dual activity (IC_50_ < 100 nM) against gametocytes and asexual blood stages, with acceptable to excellent selectivity indices (10 to > 3,000) towards HepG2 cells. Importantly, a considerable subset of these hit compounds are inhibitors of protein translation. This finding is particularly interesting in the context of a recent preprint from the Delves laboratory, showing by nascent proteomics that mature stage V gametocytes actively translate over 25% of the proteome and are thus highly translationally active (Alves *et al*., 2026). For the three potent dual-active compounds scrutinized in our study (CR-1-31-B, ONX-0914, brusatol), in vitro resistance selection coupled to WGS and drug-target interaction modelling revealed that single point mutations in the genes encoding their identified targets (or target-modifying enzyme in the case of brusatol) is sufficient to confer resistance. While this outcome is undesired from a drug development perspective, our results still identified PfeIF4A and the ribosomal A-site, and validated the proteasomal β5 subunit, as promising malaria transmission-blocking drug targets. Furthermore, clinically advanced analogs of ONX-0914 (KZR-616; Kezar Life Sciences) and CR-1-31-B (Zotatifin (eFT226); eFFECTOR Therapeutics/SJP Biotec) have undergone phase I and II clinical trials for treating autoimmune diseases, tumor malignancies and COVID-19, which may offer opportunities for drug repurposing (Johnson *et al*., 2018; Ernst *et al*., 2020; Furie *et al*., 2021; Kirk *et al*., 2021; Meric-Bernstam *et al*., 2022; Rosen *et al*., 2023)

## MATERIALS AND METHODS

### Plasmodium falciparum in vitro culture

All parasite lines were cultured in human erythrocytes (Blutspende SRK Zürich, blood groups AB+ or B+) in RPMI 1640 medium (10.44 g/l) supplemented with 2 mM choline chloride, 370 μM hypoxanthine, 100 μg/ml neomycin, 0.5% Albumax II and buffered by 24 mM sodium bicarbonate and 25 mM HEPES. To suppress sexual commitment in NF54/iGP1 and NF54/iGP1_RE9H^Pfs16^ parasites, the medium was further supplemented with 2.5 mM D-(+)-glucosamine hydrochloride (GlcN), ensuring activity of the *glmS* ribozyme. Synchronization of asexual parasites was performed using consecutive sorbitol treatments (Lambros and Vanderberg, 1979). Cultures were incubated at 37 °C in air-tight incubation chambers containing 3% O_2_, 4% CO_2_, and 93% N_2_.

### Cloning of transfection vectors

The pHF_gC-cg6#3 CRISPR/Cas9 plasmid was generated by T4 DNA ligase-dependent insertion of annealed complementary oligonucleotides (cg6#3_sg1F, cg6#3_sg1R) encoding the single guide RNA (sgRNA) target sequence sgt_cg6#3 along with compatible single-stranded overhangs into BsaI-digested pHF_gC(Filarsky *et al*., 2018). The sgt_cg6#3 target sequence (ctttaaaaatttattcgaactgg) is positioned 97-119 bp upstream of the *cg6* STOP codon and has been designed using CHOPC(Labun *et al*., 2016). The pD_cg6_pfs16-RE9H donor plasmid was generated by Gibson assembly(Gibson *et al*., 2009) of six PCR fragments. PCR fragment 1 (540 bp) represents the 5’ homology box (HB) (covering the last 166 bp of the PbDT terminator region at the 3’ end of the GDV1 overexpression cassette previously inserted into the *cg6* locus (Boltryk *et al*., 2021) and the adjacent *cg6* coding region spanning 425-100 bp upstream of the STOP codon) and was amplified from NF54 wild type gDNA using primers gib1_F and gib4_R. PCR fragment 2 (1,331 bp) corresponds to the *pfs16* upstream promoter region amplified from NF54 wild type gDNA using primers gib3_F and gib6 R. PCR fragment 3 (1,692 bp) corresponds to the *re9h* coding sequence amplified from plasmid pD_ulg8_r(Brancucci *et al*., 2025) using primers gib5F and gib8_R. PCR fragment 4 (616 bp) corresponds to the *hrp2* terminator region amplified from plasmid pBF_gC_cg6 (Boltryk *et al*., 2021). PCR fragment 5 (540 bp) represents the 3’ HB (covering the last 54 bp of the *cg6* coding sequence until 435 bp downstream of the STOP codon) and was amplified from NF54 wild type gDNA using primers gib9_F and gib12_R. PCR fragment 6 represented the pD donor plasmid backbone amplified from pUC19 using primers gib11_F and gib2_R. Oligonucleotide sequences used are provided in Supplementary Table 5B.

### Transfection and selection of transgenic NF54/iGP1^Pfs16^ parasites

250 µl of packed red blood cells containing young NF54/iGP1 ring stage parasites (parasitemia between 5% and 10%) were co-transfected as described previously (Filarsky *et al*., 2018). In brief, 50 µg each of the pHF_gC-cg6#3 CRISPR/Cas9 and pD_cg6_pfs16-RE9H donor plasmid were transfected using a Bio-Rad Gene Pulser Xcell electroporation system and 2 mm Bio-Rad cuvettes (single exponential pulse at 310V, 250 µF). 24 hours post transfection, parasites were exposed to 5 nM WR99210 for six days with daily medium changes. Subsequently, the medium was changed every second day until stable parasite propagation was observed. Successful insertion of the RE9H expression cassette directly downstream of the previously inserted GDV1 overexpression cassette (Boltryk *et al*., 2021) as well as the absence of unedited parasites in the transgenic population was confirmed by PCR on genomic DNA. Oligonucleotide sequences are provided in Table S5.

### Induction of sexual commitment and gametocyte cultures

Sexual commitment in NF54/iGP1_RE9H^Pfs16^ parasites was induced by removing GlcN and adding 1.25 µM Shield-1 to synchronous ring stage cultures to trigger GDV1-GFP-DD overexpression. Sexual commitment in the Dd2-B2 line was induced by incubating parasites at 20-24 hpi in choline chloride-depleted culture medium for 24 hours (Venugopal *et al*., 2026). Following iRBC rupture and merozoite invasion, the ring stage progeny were exposed to 50 mM N-acetyl-glucosamine (GlcNAc) for six days to eliminate asexual parasites (Ponnudurai *et al*., 1986; Fivelman *et al*., 2007), followed by further culture maintenance in regular medium until completion of gametocyte maturation. Medium was changed daily for the first six days of gametocyte development and subsequently every second day. Prior to starting drug assays, gametocytemia was quantified and gametocyte morphology inspected using standard bright-field microscopy on Hemacolor-stained thin blood smears.

### Chemical compounds

Methylene blue (MB) (Sigma-Aldrich; #M9140) and chloroquine (CQ) (Sigma-Aldrich; #C6628) were used as control compounds. Compounds tested in the primary, validation and toxicity screens derived from Novartis chemical libraries of non-proprietary compounds. Dose-response activity of selected library compounds was confirmed in the 96-well plate format for halofuginone (MedChemExpress, #HY-N1584), CR-1-31-B (MedChemExpress, #HY-136453), rocaglamide A (MedChemExpress, #HY-19356), ONX-0914 (MedChemExpress, #HY-13207), cpd 390 (Mcule, Inc., #MCULE-8094100691), cpd 464 (Mcule, Inc., #MCULE-5535790633), brusatol (Merck, #SML1868) and homoharringtonine (MedChemExpress, #HY-14944).

### RE9H luciferase-based gametocyte viability assay (96-well format)

Gametocytocidal assays using the 96-well plate format were performed using stage V gametocytes (day 12 of gametocyte development) as described previously (Brancucci *et al*., 2025). In brief, 100 µl culture aliquots at a gametocytemia of 0.3% and a hematocrit of 1% (approx. 30,000 gametocytes) were transferred to the wells of 96-well cell culture plates (Corning; #353072) containing compounds of interest (in 100 µl culture medium containing a maximum of 1% DMSO) and incubated for 72 hours at 37 °C. RE9H-mediated bioluminescence was measured by transferring 90 µl of each suspension to individual wells of a black-wall 96-well plate (Greiner CELLSTAR; #7.655.086) preloaded with 10 µl D-luciferin in PBS (3.75 mg/ml) (PerkinElmer; #122799). After 5 min incubation, luminescence was quantified using either an IVIS Lumina II *in vivo* imaging system (Caliper Life Sciences, PerkinElmer), or a luminescence plate reader (TECAN SPARK, Tecan Group AG). Assay quality was monitored by calculating Z’ scores from wells of treated (50 µM MB) and untreated (1% DMSO vehicle) controls, with a quality inclusion threshold of Z’ > 0.6; Z’ score was calculated as follows: Z’ = 1 − ((3 ∗ ∂DMSO + MB)/(।μDMSO − μMB।)); *∂* = standard deviation; *μ* = mean; DMSO = untreated controls; MB = treated controls.

Data was analysed in GraphPad Prism (version 8.2.1) following averaging of luminescence counts from technical replicates. IC_50_ values were calculated using nonlinear, four parameter (variable slope) curve fitting.

### RE9H luciferase-based gametocyte viability assay (1536-well format)

Gametocytocidal assays using the 1536-well plate format were performed as indicated for the 96-well plate format, with following differences: 7.5 µl of a gametocyte culture at a gametocytemia of 0.3% and a hematocrit of 1% (approx. 2,000 gametocytes) were transferred to the wells of 1536-well black-wall cell culture plates equipped with a humidity chamber surrounding the wells (Greiner Bio-one 792091-191, custom-made for Novartis Pharma AG, black plate with clear bottom) using a 64-syringe microplate dispenser (AquaMax DW4 microplate dispenser, Molecular Devices) at a pressure of 450 millibar. Plates were spotted with 7.5 nl of compounds (in 90% DMSO) using an acoustic liquid handler (Echo 550, Beckman Coulter) prior to adding the gametocyte culture. After dispensing, the humidity chamber was filled with culture medium and plates were immediately placed into a humidified CO_2_ incubator (5% CO_2_, 19% O_2_, 76% N_2_) and incubated for 72 hours at 37 °C. RE9H-mediated bioluminescence was measured after transferring 2.5 µl of D-luciferin in PBS (3.75 mg/ml, PerkinElmer; #122799) directly to each well at room temperature (AquaMax DW4, Molecular Devices) using a pressure of 550 millibar and incubation for 5 to 15 min. Luminescence was subsequently quantified using a camera-based plate reader (Luminescence Plate Reader, LPR, GNF Systems, serial # S1792A). Assay quality was monitored by calculating Z’ scores from 32 technical replicate wells of each treated (50 µM MB) and untreated (DMSO vehicle) controls per plate. Parasites incubated with MB at 10 µM and 800 nM were included as a further reference but not used for downstream analyses. All plates of the primary and validation screens passed the Z’ inclusion threshold of 0.6.

### Processing of primary screening data

Data was analyzed using the Helios platform (Novartis Institutes of Biomedical Research), a fully automated and modular software environment for plate-based data processing, artifact correction, quality control, and dose-response analysis (Gubler *et al*., 2018). Raw image data (x) was subject to control-based normalization using the mean signals of neutral control wells (NC, untreated) and active control wells (AC, treated with 50 µM MB) and activity values calculated according to the following formula: -100(x-NC)/(AC-NC) [%]. The means of the NC and AC control wells define no activity (0%) and full inhibition (-100%), respectively. Systematic spatial artifacts were corrected using the robust local regression (RLOCREG) method, in which intensities were modeled as a smooth function of plate coordinates using locally weighted polynomial regression (Gubler *et al*., 2018).

### Validation screen and toxicity counter screen (1536-well format)

The validation screen and toxicity counter screen were performed using the 1536-well plate format with 8-point 1:4 dilution series covering a compound concentration range of 5 µM to 0.3 nM. Stage V gametocytes were used in the validation screen as described above, using technical quadruplicates. Toxicity counter screening was performed with HepG2 cells (7.5 µl per well of 40,000 cells/ml) in combination with the CellTiter-Glo 2.0 viability assay reagent (2.5 µl per well) (Promega Corporation, USA) in technical quadruplicates. Luminescence of the validation screen was quantified as indicated above, while toxicity levels were measured using a Pherastar FS BMG Labtech, # 470-084. Structural clustering was performed using the Butina Clustering function in RDKit [RDKit: Open-source cheminformatics; http://www.rdkit.org] with morgan fingerprints (radius 3, 2048 bits), and a cutoff distance of 0.35.

### Asexual blood stage viability assay

Antimalarial activity against asexual blood-stage parasites was assessed using a SYBR Green I-based proliferation assay [[PMID:17371812](https://pubmed.ncbi.nlm.nih.gov/17371812/)]. Parasites were cultured in human O⁺ erythrocytes in RPMI-1640 medium supplemented with 0.5% Albumax II, 200 µM hypoxanthine, 50 mg/l gentamicin sulphate, 35 mM HEPES, 2.0 g/l sodium bicarbonate, and 11 mM glucose, and maintained at 2–3% hematocrit with parasitemia ranging from 1–8% at 37 °C in 5% CO2, following standard continuous culture protocols [[PMID:781840](https://pubmed.ncbi.nlm.nih.gov/781840/)]. For susceptibility testing, unsynchronized infected erythrocytes were adjusted to 0.3% parasitemia and 2.5% hematocrit. Test compounds, serially diluted 3-fold in DMSO across ten concentrations (final DMSO concentration ≤0.09% v/v), were dispensed into 384-well plates in duplicate, followed by 50 µl of the parasite suspension per well. Plates were incubated for 72 hours at 37 °C in 5% CO_2_. Parasite DNA content was then quantified by adding lysis buffer containing 5 mM EDTA, 1.6% Triton X-100, 20 mM Tris-HCl, 0.16% saponin, and 0.1% SYBR Green I, followed by a 24-hour incubation at room temperature protected from light. Fluorescence (excitation 483 nm, emission 530 nm) was measured using a plate reader (BMG LabTech CLARIOstar). Percent activity was calculated relative to a neutral control (DMSO alone, defined as 0% inhibition) and an active control (10 µM mefloquine, defined as 100% inhibition), and dose-response curves were fitted to a four-parameter logistic function to derive IC_50_ values.

Dose-response activities of selected hit compounds on asexual parasites was confirmed using a [^3^H]hypoxanthine incorporation assay as described previously (Snyder *et al*., 2007). In brief, asexual parasite cultures (200 µl, 0.3% parasitemia, 1.25% hematocrit) were incubated with compounds in hypoxanthine-free medium in 96-well plates (Corning; #353072). After 48 hours, [^3^H]hypoxanthine was added to all wells (0.25 μCi/well) and plates were incubated for an additional 24 hours. Parasites were harvested using a Microbeta FilterMate cell harvester (Perkin Elmer, Waltham, USA) and radioactivity was quantified using a MicroBeta2 liquid scintillation counter (Perkin Elmer, Waltham, USA). Following subtraction of the signal emitted from uninfected erythrocytes and normalization to untreated controls, the data was analysed GraphPad Prism (version 8.2.1) and IC_50_ values were calculated using nonlinear, four parameter (variable slope) curve fitting.

### Gamete formation assay

Sexual commitment was induced in NF54/iGP1_RE9H^pfs16^ parasites as described above. Gametocytes were subsequently cultured in medium containing 10% heat-inactivated human serum (instead of 0.5% Albumax II). Male and female gamete activation assays were performed using mature gametocytes according to established protocols(Delves *et al*., 2016). In brief, gamete formation was quantified following incubation of 2 μl packed RBCs containing stage V gametocytes (day 13) in 50 µl activation medium (serum medium supplemented with 100 µM xanthurenic acid). To quantify male gamete formation, parasites were immediately transferred to a Neubauer chamber and incubated in a humid chamber at room temperature for 15 minutes. Exflagellation rates were determined by microscopy of four grids of a Neubauer chamber (4 mm^2^ ≙ 400 nl) at a 400x magnification (DM1000, Leica Microsystems) using the following formula 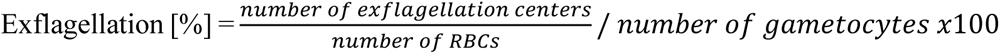. To quantify female gamete formation, mouse anti-Pfs25 primary antibodies (MR4-28, lot 70013640; 1:2000) and goat anti-mouse secondary antibodies coupled to Alexa Fluor 594 (Thermo Fisher, #A-11005; 1:250) were added to the activation medium.

Parasites were incubated in the dark rotating at room temperature overnight. Following staining with 1x SYBR Green I DNA dye (Invitrogen #S7563, diluted 1:10,000), samples were diluted to a hematocrit of 0.1% and 200 µl were transferred to a high content imaging plate (Greiner CELLCOAT microplate 655948, Poly-D-Lysine, flat μClear bottom). Cells were subsequently allowed to settle for 15-20 min and 36 images were taken per condition in three channels (SYBR Green I (Ex.: 472/30; Em.:520/35); Alexa Fluor 594 (Ex.: 562/40: Em.:624/40); transmitted light) using a Plan-Apochromat 40x objective (Molecular Devices, #1-6300-0297) in conjunction with an ImageXpress Micro widefield high content screening system (Molecular Devices). The proportion of SYBR Green I/Alexa Fluor 594-positive female gametocytes among all SYBR Green I-positive cells was quantified using an automated image analysis workflow created using the MetaXpress software (version 6.5.4.532, Molecular Devices), as described previously (Kuehnel *et al*., 2023).

### Mosquito feeding assays

Standard membrane feeding assays (SMFAs) were performed by TropIQ Health Sciences (Nijmegen, The Netherlands) according to standard procedures (Vos *et al*., 2015; Dechering *et al*., 2017). Test compounds in DMSO were dissolved in culture medium to achieve final concentrations equivalent to the respective IC_50_ or 10x IC_50_ determined in the *in vitro* gametocytocidal dose-response assays. Concentration of the DMSO vehicle was kept constant at 0.1%. Stage V gametocytes of *P. falciparum* strain NF54-HGL were incubated in the presence of compounds for 48 hours at 37 °C. Gametocyte cultures were subsequently adjusted to a hematocrit of 50% in human serum and the sample fed to two-day old, starved *A. stephensi* mosquitoes. Mosquito infection was analyzed eight days post-feeding by quantifying luminescence emitted from whole mosquitoes as described previously (Vos *et al*., 2015). Gametocytes exposed to the DMSO vehicle (0.1%) and dihydroartemisinin (10 μM) served as negative and positive controls, respectively. Samples were tested in duplicate experiments using 12 (negative control) or 24 mosquitoes (positive control and test compounds) each per feed. Mosquitoes were considered infected when the luminescence of test mosquitoes increased by at least five standard deviations from the mean background signal measured from the 24 non-infected control mosquitoes. The following quality thresholds were applied: >50% infection prevalence in the negative controls; >70% reduction in oocyst intensity in the positive control.

### Resistance selection and sensitivity testing

90 ml of Dd2-B2 asexual blood stage parasites synchronized to a time window of 0-8 hpi at a 5% hematocrit and a 4-5% parasitemia were exposed to drug concentrations equivalent to 3x the IC_50_ (brusatol, CR-1-31-B) or 6x the IC_50_ (ONX-0914) against asexual NF54 wild type parasites. Parasite morphology was monitored regularly by visual inspection of Hemacolor-stained thin blood smears during the first days following initiation of treatment to confirm drug-mediated killing. Medium containing compounds was changed every 24 hours during the first eight days and subsequently every 48 hours. Cultures were maintained for up to two months, with continuous monitoring via thin blood smears to detect the emergence of resistant parasites. After selection of stably propagating resistant parasite populations, clonal lines were established using limiting dilution(Thomas *et al*., 2016).

Drug sensitivity testing of asexual parasites against ONX-0914 and brusatol was performed using synchronized young ring stage parasites at 0.5% parasitemia and 1.25% hematocrit. Parasites were exposed to compounds in 96-well plates (Corning, 96-well cell culture plate, flat bottom, # 3596) for 48 hrs. Parasitemia was determined at the start and end of the assay after incubation with 1x SYBR Green I DNA stain (1:10,000) (Invitrogen #S7563) and a flow cytometry-based readout (MACSQuant VYB, Miltenyi Biotec) (Fig. S8). CR-1-31-B was tested using the [^3^H]hypoxanthine incorporation assay as explained above. For drug sensitivity testing on stage V gametocytes, gametocyte cultures were prepared according to the culture protocol explained above. Mature stage V gametocytes (day 12) at 0.5-2% gametocytemia and 2% hematocrit were exposed to compounds in 96-well plates. After an incubation period of 72 hrs, 50 μl of gametocyte suspension from each well was transferred to a new 96-well plate preloaded with 50 μl culture medium containing 400 nM MitoTracker Red CMXRos (Invitrogen;# M46752) and 1:5,000 SYBR Green I (Invitrogen #S7563) and stained for 20 min at 37 °C. Stained cell suspensions were further diluted to 0.1% hematocrit in 50 μl culture medium and transferred to high content imaging plates and imaged using an ImageXpress Micro widefield high content screening system (Molecular Devices) as described above. Viable gametocytes were quantified as the proportion of SYBR Green I/MitoTracker-positive gametocytes among all SYBR Green I-positive cells based on MitoTracker signal intensity and cellular shape using the MetaXpress software as described previously (Brancucci *et al*., 2025).

### Whole genome sequencing and variant analysis

500 μl of packed RBCs at a parasitemia of 4-6% were resuspended in 1 ml saponin solution (0.15% in 1x PBS) and incubated on ice for 10 minutes to selectively lyse RBCs. Released parasites were pelleted (5 min, 1,500 g) and washed twice in 1x PBS. Parasite gDNA was extracted from the parasite pellets using the QIAamp DNA Blood kit mini (Qiagen, # 51104) according to the manufacturer’s instructions. To avoid amplification bias of AT-rich genomic sequences, library preparation was performed without PCR amplification using the KAPA HyperPrep Kit (Roche). Libraries were subsequently sequenced on the NextSeq 500 system using the NextSeq 500 Mid Output Kit (Illumina) and a read-length of 2x150 bp (paired-end sequencing), aiming at a 50-100x coverage. WGS reads were analyzed as previously described (Hitz *et al*., 2021). In brief, raw reads were mapped to the *P*. *falciparum* 3D7 reference genome (PlasmoDB v68) using the Burrows Wheeler Aligner (version 0.7.18) with default parameters. BAM files were sorted by coordinate, indexed, and read groups assigned using Picard (version 3.2.0). Single nucleotide polymorphisms (SNPs) and insertions/deletions (Indels) were identified with the HaplotypeCaller tool from the Genome Analysis Toolkit (GATK, version 4.6.0.0) in GVCF mode to enable multisample analysis. The resulting GVCF files from all samples within a population were merged and genotyped. To assess the functional impact of the identified variants, annotation was performed using SnpEff (version 5.1). The variants were filtered by (i) variant quality score (QUAL > 500) to confirm variation at the observed site; (ii) genotype quality of the wild type parent sample (GQ = 99); (iii) sites at which the genotype of the two wild type parent samples correspond to the reference genome. Variants that passed these criteria were verified visually by comparing reads of all samples using the Integrative Genomics Viewer (version 2.5.0). In the case of the brusatol-resistant population C, a CNV analysis was performed in addition to variant analysis. CNV was estimated by comparing the number of reads (RPKM) for a gene between the parental samples and resistant clones using Artemis (version 16.0.17) (Milne *et al*., 2022).

### Oxford Nanopore sequencing of rRNA

500 μl of packed RBCs at a parasitemia of 4-6% were resuspended in 1 ml saponin solution (0.15% in 1x PBS) and incubated on ice for 10 minutes. Parasites were pelleted (5 min, 1,500 g) and washed twice in 1x PBS. The parasite pellet was suspended in Trizol (Invitrogen, # 15596026) and total RNA extracted by ethanol precipitation. The resulting total RNA was then poly-A-tailed using *E. coli* Poly(A) Polymerase (NEB, # M0276) and purified with the Agencourt RNAClean XP beads (Beckman Coulter, # A63987) and isopropanol precipitation. RNA quality was assessed on an Agilent TapeStation (Agilent, 4150 TapeStation system).

Sequencing libraries were prepared using the Direct RNA Sequencing Kit (Oxford Nanopore Technologies, # SQK-RNA004) following the manufacturer’s instructions with 700 ng of total RNA input and using the Induro Reverse Transcriptase (NEB, # M0681) for reverse transcription. Between 96 to 180 ng were loaded onto an RNA flow cell (Oxford Nanopore Technologies, # FLO-MIN004RA) depending on the samples, and the sequencing ran for 1.5 to 2 hrs.

As a non-modified, negative control, *in vitro* transcribed RNA of the A-type ribosomes was sequenced in parallel. In brief, two overlapping fragments covering the sequence of the A-type (PF3D7_0726000) 28S rRNA were amplified from wild-type *P. falciparum* (strain NF54) genomic DNA by PCR. Poly-A tails (30 A’s) for subsequent *in vitro* transcription (IVT) were added by including a poly-T tail in the reverse primer of each fragment. PCR primer sequences can be found in Table S5. The IVT was performed using the MEGAscript Kit (Invitrogen, # AM1334) according to the manufacturer’s instructions and the resulting RNA was purified with the Zymo Kit Clean and Concentrator kit (Zymo Research, # R1013).

Raw pod5 files were first merged using pod5 ‘merge’ (version 0.3.23, https://github.com/nanoporetech/pod5-file-format) and base-called using dorado ‘basecaller’ (version 0.7.4, https://github.com/nanoporetech/dorado), with basecalling model rna004_130bps_sup@v5.0.0 and options ‘—emit-moves –emit-sam’. Non primary and supplementary alignments were removed using samtools (Danecek *et al*., 2021) ‘view’ (option ‘-F 2304’) (version 1.15.1), sorted using samtools ‘sort’, and only chromosome 7 (Pf3D7_07_v3) encoding for the A-type rDNA was retained. The resulting BAM files were subsequently processed using the *align* command of Uncalled4^2^ (version 4.1.0) (Kovaka, 2025).

To calculate the current values for every position of each read in the native and *in vitro* control RNA, a tsv output file was generated using Uncalled4 ‘align’ with option ‘--tsv-out’. The values corresponding to the positions of interest were extracted and visualized using ggplot (Wickham, 2016) in RStudio (R Core Team, 2024) (version 2024.09.0+375). To get a visual overview of the current deviations from the model, heatmaps were generated using uncalled4 ‘*trackplot’*, targeting a region comprising a hundred nucleotide upstream and downstream of the position of interest.

### RNA extraction and Nm-VAQ

160 μl of packed RBCs at a parasitemia of 4-6% were resuspended in 1 ml saponin solution (0.15% in 1x PBS) and incubated on ice for 10 minutes. Parasites were pelleted (5 min, 1,500 g) and washed twice in 1x PBS. The pellet was resuspended in 1 ml TRIzol (Thermo Fisher Scientific) and incubated at 37°C for 5 min. 200 μl chloroform (Sigma Aldrich) were added and the sample spun down at 20,000 g for 15 min at 4 °C. 500 μl of the aqueous phase were transferred into a fresh Eppendorf tube, from which RNA was extracted using the Zymo Direct-zol RNA MicroPrep kit (Zymo Research) according to the manufacturer’s instructions.

To detect site-specific rRNA base 2’-O-methylation, we applied the Nm-VAQ method (Tang, 2024). Specifically, 200 ng of parasite RNA were hybridized separately with 10 pmol, 3.33 pmol, 1.11 pmol or 0.37 pmol RNA/DNA chimera (Genscript). Subsequently, the hybridized RNA/chimera mixture was split and half was subjected to RNase H digestion. Next, the primers 3’ of the suspected methylated nucleotide (Nm) site (RT primer 1 for rRNA bases 2665-2667 and 2691, RT primer 2 for rRNA base 3281) were annealed to the samples and reverse transcription performed using the Invitrogen SuperScript IV kit and M-MLV Reverse Transcriptase (Promega). The resulting cDNA was amplified using two forward primers and one reverse primer; one pair resulted in the amplification of the sequence containing the Nm site (longer sequence) and the other amplified the section downstream of the Nm site (shorter sequence). The amplified DNA was stained with peqGreen (VWR) and loaded on a 2% agarose gel, run at 80V for 40 min and imaged using a VisionCapt Imager (Vilber). Sequences of primers and chimera are provided in Table S5.

### Drug-target interaction modelling

#### Complex preparation

The homology model of PfeIF4A1 (Q8IKF0_PLAF7) was generated employing SWISS-MODEL (Waterhouse *et al*., 2018a), using the X-ray structure of human eIF4A1 bound to ANP (phosphoaminophosphonic acid-adenylate ester), RNA and silvestrol (PDB ID: 9 (Naineni *et al*., 2024)) as a template. The Mg^2+^ ion, ANP cofactor, silvestrol, and RNA from 9avr were subsequently transferred to the homology model. The ANP cofactor was converted to ATP, its physiologically relevant form, by replacing the nitrogen atom in the aminophosphonate group with an oxygen atom. The ATP protonation state was adjusted to pH 7.4 to account for the surrounding hydrogen bond network and interactions with the Mg^2+^ ion. Hydrogen atoms were added to the protein and RNA using REDUCE (Word *et al*., 1999). For the Thr100Ala and Gln185Lys mutants, side chains were modeled using the mutagenesis wizard tool in PyMOL (DeLano, W.L., 2002).

For the Pf20S proteasome subunit β5 (PF3D7_1011400), the cryo-EM structure of the Pf20S in complex with the WLW-vs inhibitor covalently bound to the catalytic Thr1 (PDB ID: 8g6f(Hsu and Li, 2023) was used as a template. The resistance mutation Ala117Asp in β5 was manually reverted back to the wild type form using PyMOL. To make MD calculations more tractable, only subunits β5 (auth chain L) and β6 (auth chain M), which interact directly with the WLW-vs inhibitor, were retained. Hydrogen atoms were added using REDUCE, and the Met105Ile mutation was introduced using PyMOL.

For PF3D7_1354300, annotated as a putative rRNA methyltransferase, a homology model was built with SWISS-MODEL (Waterhouse *et al*., 2018b), using the cryo-EM structure of the 27S pre-rRNA (guanosine(2922)-2’-O)- methyltransferase SPB1 from *S. cerevisiae* (PDB ID: 7nac (Cruz *et al*., 2022) as a template.

#### Constrained docking protocol

The CR-1-31-B and ONX-0914 inhibitors were positioned using experimentally resolved inhibitors as templates. For ONX-0914, the WLW-vs inhibitor from the Pf20S assembly (PDB ID: 8g6f, 7F1, auth chain L) served as a template, while the silvestrol inhibitor from the human eIF4A1 complexed with ANP, RNA and silvestrol (PDB ID: 9avr, A1AG8, auth chain B) was used for CR-1-31-B. RDKit (Landrum, no date) was employed to identify the maximum common substructure between each inhibitor and its corresponding template, which was then used as a constraint to generate conformations. A total of 10 conformations were generated per inhibitor. Each conformation was locally minimized in the presence of the receptor using AutoDock Vina(Trott and Olson, 2010), and the top-scoring conformation was selected. Both inhibitors and receptors were processed and converted to PDBQT format using Meeko(Santos-Martins *et al*., 2025). Docking boxes were centered on the template ligand positions, with dimensions set to 20 × 20 × 20 Å. The resulting PDBQT files were subsequently converted back to MOL format using Meeko for downstream calculations.

#### Molecular Dynamics simulation protocol

To investigate the effects of resistance mutations on inhibitor binding, unbiased MD simulations were performed on the various receptor–inhibitor complexes. All systems underwent identical preparation and simulation protocols. Simulations were carried out using OpenMM 8.2 (Eastman *et al*., 2017), employing the ff19SB force field (Tian *et al*., 2020) for proteins, OL3 (Zgarbová *et al*., 2011) for RNA (when present), and GAFF2.11 (Wang *et al*., 2004) for inhibitors and cofactors. Parameters for small molecules were generated using antechamber (Wang *et al*., 2004). Charged N- and C-terminal patches were applied following the standard Amber protocol. ONX-0914 was simulated in its pre-reactive state, without covalent bonding to Thr1. A 12 Å cutoff was used for non-bonded interactions, and long-range electrostatics were treated with the Particle Mesh Ewald (PME) method(Petersen, 1995), using mixed precision to maintain numerical accuracy. The SHAKE algorithm (Weeks, Chandler and Andersen, 1971) was applied to constrain bonds involving hydrogen atoms, while SETTLE was used for water molecules (Miyamoto and Kollman, 1992).

To prevent artifacts from slow water diffusion in enclosed pockets, particularly around inhibitors or cofactors (e.g., ATP and Mg^2+^ in the eIF4A1–RNA complex), the first hydration shell was predicted using WaterKit (Eberhardt and Forli, 2023). Hydration site prediction was performed in the presence of inhibitors and cofactors, following WaterKit’s standard procedure. A single set of water molecule positions was generated using a box centered on the complex, sized to encompass the entire complex. Default parameters were used (temperature: 300 K; number of layers: 3). The receptor–inhibitor complex and predicted hydration shell were then placed in an orthorhombic box, with 12 Å padding from the nearest atom, and solvated with explicit TIP3P water molecules. The system’s net charge was neutralized with K⁺/Cl⁻ counterions using tleap from the Amber package (Case *et al*., 2023).

Prior to production simulations, steric clashes were resolved through energy minimization without restraints. This was followed by a 3 ns equilibration under NPT condition (1 atm, 300 K) using a Monte Carlo barostat and Langevin middle integrator (“OpenMM Integrator,” no date), with a 2 fs time step. During equilibration, positional restraints were applied to all heavy atoms except water molecules and were linearly reduced every 500 ps, from 2.5 kcal/mol/Å² to 0 kcal/mol/Å² in the final 500 ps. Production runs were performed in NVT condition at 300 K, using the Langevin middle integrator and hydrogen mass repartitioning (HMR) to 4 amu, allowing a 4 fs integration time step.

A total of 10 replicates of 150 ns were generated for wild-type and mutant Met105Ile Pf20S β5/β6 subunits with bound ONX-0914, and for wild-type and mutant Thr100Ala PfeIF4A1–RNA with bound CR-1-31-B. For wild-type and mutant Gln185Lys PfeIF4A1–RNA with bound CR-1-31-B, unbiased MD simulations were halted after the equilibration, and funnel metadynamics simulations were pursued. System conformations were saved every 1 ps for subsequent analysis.

#### Funnel metadynamics protocol

Funnel metadynamics (Fun-metaD) simulations were carried out to estimate the absolute binding free energy (ABFE) profiles of the CR-1-31-B inhibitor in complex with wild-type and Gln185Lys mutant PfeIF4A1–RNA. Fun-metaD was performed using the BioSimSpace (BSS) implementation (Hedges *et al*., 2019), in combination with OpenMM 8.2. The final snapshot from the equilibration phase served as the starting point for the simulations, which were run under NVT conditions using the Langevin middle integrator at 300 K, with hydrogen mass repartitioning (HMR) set to 4 amu and a 4 fs integration time step.

A combination of Well-Tempered (WT) metadynamics and funnel-shaped restraints was used. A history-dependent bias was applied along two collective variables (CVs): projection and extent. The first CV measures the distance along the axis-vector defined by the coordinates of the points P_0_ and P_1_, automatically defined with BSS based on the inhibitor and PfeIF4A1 coordinates. The second CV measures the distance along the second axis-vector orthogonal to the first axis-vector. The funnel width was set to 6 Å, with lower and upper bounds at 0 Å and 45 Å, respectively, and an inflection point at 25 Å. The funnel geometry was visually validated to ensure that P0 and P1 provided an unobstructed unbinding path and prevented the inhibitor from sampling non-relevant regions. Gaussian hills with an initial height of 0.35 kcal/mol were deposited every 1 ps. These hills were (Michaud-Agrawal *et al*., 2011; Gowers *et al*., 2016)scaled using the WT scheme with a bias factor of 10. The hill widths for the projection and extent CVs were set to 0.25 Å and 0.5 Å, respectively.

In total, 10 replicates of 100 ns were generated for both the wild-type and Gln185Lys mutant PfeIF4A1–RNA + CR-1-31-B systems. ABFE values were computed using PLUMED (Tribello *et al*., 2014), with corrections applied to account for the loss of translational and rotational entropy of the unbound inhibitor imposed by the funnel-shaped restraint.

#### Relative free-energy calculations

Mutation-induced changes in ONX-0914 binding affinity were estimated using alchemical free energy perturbation (FEP) with the FEP/REST protocol implemented in Desmond (Schrödinger Release 2025-4: Desmond Molecular Dynamics System, D. E. Shaw Research, New York, NY, 2024. Maestro-Desmond Interoperability Tools, Schrödinger, New York, NY, 2025). Simulations were based on the experimentally determined protein structure (PDB ID: 8G6E) and employed the OPLS4 force field for protein and ligand. Ligand torsional parameters were refined using the Schrödinger Force Field Builder. The Met105Ile mutation was modeled as a single-residue alchemical transformation in the presence of the bound ligand. Enhanced sampling was achieved using Replica Exchange with Solute Tempering (REST) applied to the mutating residue and surrounding region. The alchemical transformation was performed using 12 λ windows, each simulated for 5 ns. Free energy differences were obtained by integrating the free energy gradients across all λ states using the standard Desmond FEP workflow to yield the relative binding free energy change (ΔΔG) associated with the mutation.

Relative binding free energy changes upon the Met105Ile mutation were also estimated using alchemical free energy perturbation (FEP) simulations performed with BioSimSpace(Hedges *et al*., 2019) interfaced with GROMACS, with GPU acceleration enabled for all major simulation steps. Wild-type and mutant proteins were parameterized using the AMBER ff14SB force field. The mutation was represented using a dual-topology alchemical transformation, generated by atom mapping between the wild-type and mutant residues within a residue-centered region of interest using the BioSimSpace.Align.matchAtoms method. Apo and holo systems were solvated in explicit TIP3P water within a rhombic dodecahedral simulation box with a minimum solute padding of 15 Å, and neutralized with Na⁺ and Cl⁻ ions to reach 0.15 M ionic strength. Following energy minimization with backbone restraints, the systems were equilibrated for 5 ns in the NVT ensemble at 300 K using a 1 fs integration time step. Hydrogen-containing bonds were constrained using LINCS, and long-range electrostatics were treated using particle mesh Ewald (PME) with a 0.8 nm cutoff for both electrostatic and Lennard–Jones interactions. The alchemical protocol used 12 λ windows, each simulated for 5 ns. Free energy differences were calculated using the multistate Bennett acceptance ratio (MBAR) method to obtain the relative binding free energy change (ΔΔG).

#### MD trajectory analysis

All analyses were performed using the MDAnalysis Python package (Michaud-Agrawal *et al*., 2011; Gowers *et al*., 2016), and receptor-inhibitor interactions were monitored using ProLIF (Bouysset and Fiorucci, 2021).

### Statistical analysis

Library screenings were performed in a single experiment (high throughput primary screen) or technical quadruplicates (high throughput validation and toxicity counter screens). The primary screen was performed using single datapoints measured from stage V gametocytes exposed to test compounds at 5 µM. Compounds resulting in relative luminescence unit (RLU) intensities 30% below the mean obtained from all negative control wells (untreated) were considered as primary hits. The dose-response validation screen and toxicity counter screen were performed using an 8-point 1:4 dilution series covering a concentration range of 5 µM to 0.3 nM. Dose-response compound screening against asexual parasites was performed using a 10-point 1:3 dilution series covering a concentration range of 10 µM to 0.5 nM. Unless stated otherwise, all other experiments were performed in biological triplicates. Data from high throughput assays (primary screen, validation screen, toxicity screen) were handled using Helios. Representative dose-response curves were fitted using GraphPad Prism (version 8.2.1) by applying a nonlinear, four parameter (variable slope) method with error bars representing the standard error of the mean. Except for dose-response experiments, error bars represent standard deviations. If not stated otherwise, significance testing was performed using unpaired, two-tailed t-tests.

## Acknowledgements

We are grateful to Thierry T. Diagana for initiating the collaboration between Biomedical Research, Novartis and Swiss TPH. The authors would also like to thank Dr. Patricia Bravo for sharing her expertise, time and WGS analysis pipeline, as well as Dr. Natalie Wiedemar for her WGS and CNV analysis code. This work was supported by grants from the Fondation Pasteur Suisse (TSV), the Swiss National Science Foundation (BSCGI0_157729, TSV; 320030-231866, NMBB), Medicines for Malaria Venture (PO17/00583, MR), Novartis Pharma AG, Basel, Switzerland (DB), a Wellcome Trust Career Development Award (325247/Z/25/Z, VT), as well as an ERC Starting Grant (947819) and baseline funding from the Institut Pasteur and INSERM (SB).

## Author contributions

Conceptualization: NMBB, TSV, MR, DB

Methodology: NMBB, TSV, MR, LS, CG, SB, ML, HB, LTA, DS

Investigation: LS, NMBB, CG, OC, BTT, AP, KS, SB, JS, VT, WAG, LP

Formal Analysis: NMBB, TSV, CG, LS, SR, JE, DIB, AHM

Visualization: NMBB, LS

Funding acquisition: TSV, NMBB, MR, VT, SB

Project administration: NMBB, TSV

Resources: NMBB, TSV, MR, DB, SB

Supervision: NMBB, TSV, MR, DB, SB, MAL

Validation: NMBB, TSV, MR, DB, VT, SB, MAL, JS

Writing – original draft: NMBB, LS, TSV

Writing – review & editing: NMBB, LS, TSV, MR, DB, HB, SR, JS, CG, VT, MAL

## Data and materials availability

All data generated in this study are present in the main manuscript or in the Supplementary Materials. Correspondence should be addressed to MR, TSV or NMBB.

