## Supplementary Information for "A high-throughput compound screen identifies multiple druggable targets in Plasmodium falciparum transmission stages"

**This Supplementary Information file includes:**

Figs. S1 to S8

Tables S1 to S5

Datasets S1 and S2

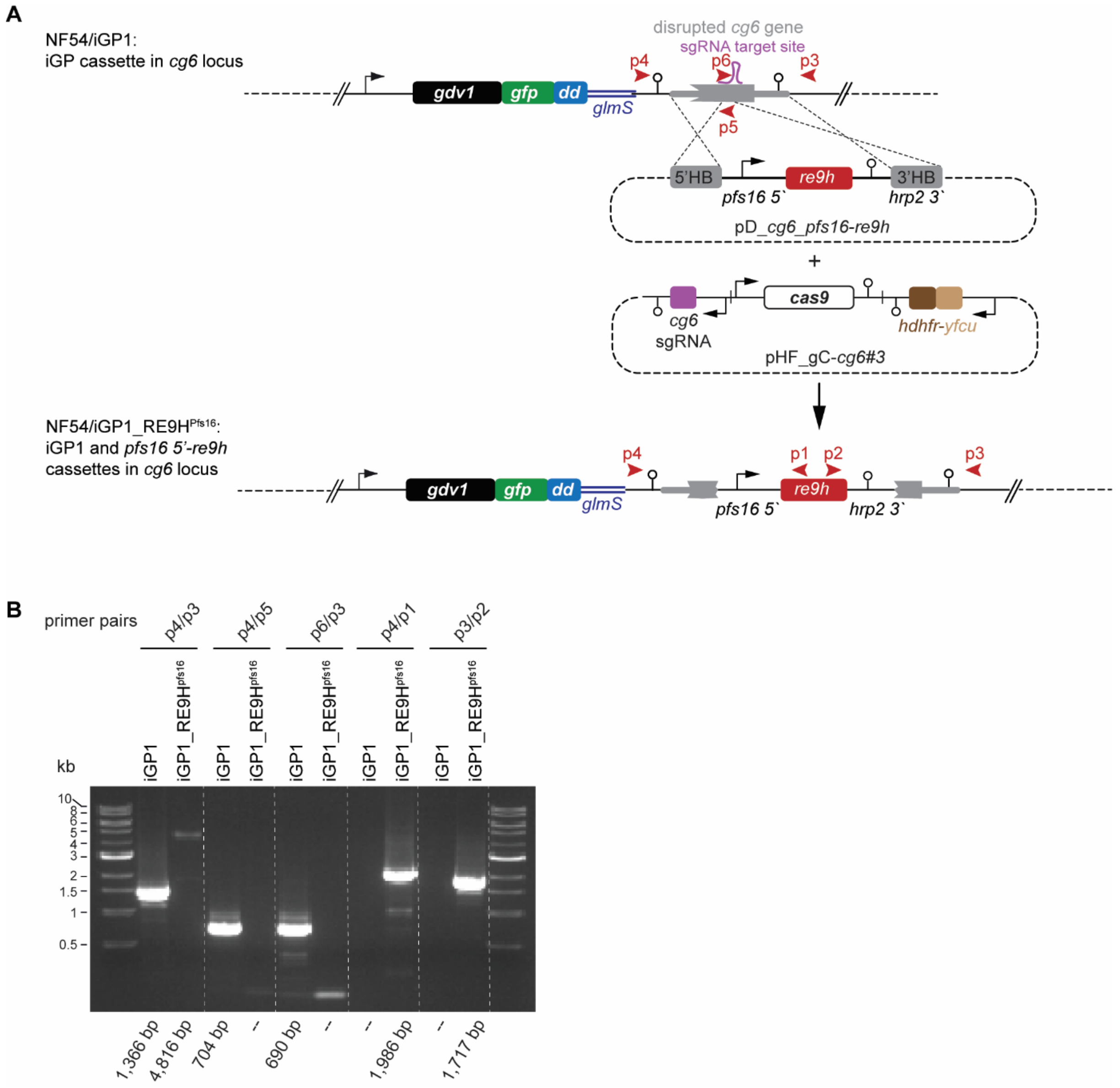

**Figure S1. Generation of the NF54/iGP1_RE9Hpfs16 gametocyte reporter line.**

**A)** Schematic map of the chromosomal *gdv1-gfp-dd-glmS* expression cassette at the disrupted *cg6* locus in NF54/iGP1 parasites (Boltryk et al., 2021) (top), the pD_cg6_pfs16-RE9H donor and pHF_gC-cg6#3 CRISPR/Cas9 plasmids (center), and the double gene-edited *cg6* locus carrying the transgene cassette for *pfs16* promoter-driven RE9H luciferase expression inserted directly downstream of the *gdv1-gfp-dd-glmS* expression cassette (bottom) in NF54/iGP1_RE9Hpfs16 parasites. The relative position of the sgRNA target site is shown in purple. The pD_cg6_pfs16-RE9H plasmid contains the *re9h* gene controlled by the *pfs16* promoter (pfs16 5') and the histidine-rich protein 2 terminator (hrp2-3'), flanked by 5' and 3' homology boxes (HB) for homology-directed repair (grey). The pHF_gC-cg6#3 plasmid contains expression cassettes for the SpCas9 nuclease (white), the sgRNA (purple) and the h*dhfr-yfcu* positive-negative drug selection marker (brown). Relative positions of PCR primer binding sites are indicated by red arrowheads. glmS, glmS riboswitch element. dd, FKBP-derived destabilization domain. **(B)** Results of diagnostic PCRs performed on gDNA from NF54/iGP1 (control) and

NF54/iGP1_RE9H[pfs16] parasites. Primer combinations used to amplify sequences specific to the *cg6* locus prior to or after insertion of the *re9h* expression cassette in NF54/iGP1 or NF54/iGP1_RE9Hpfs16 parasites, respectively, are shown on top. Expected PCR fragment sizes (bps) are indicated.

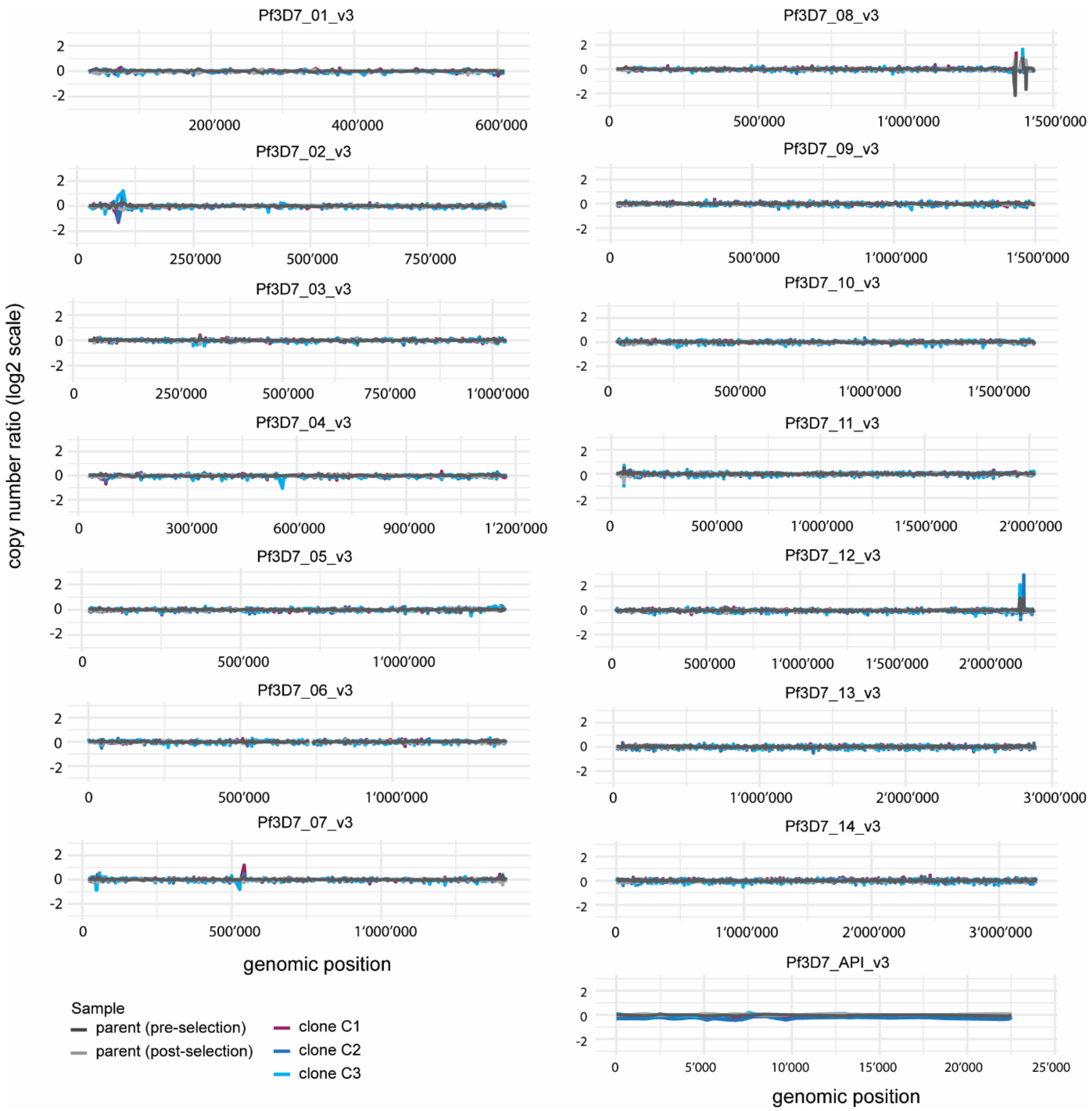

**Figure S2. CNV analysis of brusatol-resistant clones C1, C2 and C3.**

The copy number variant (CNV) ratio (log2-scaled) of wild-type parent populations collected pre- and post-selection (untreated control), brusatol-resistant clones C1 (magenta), C2 (dark blue) and C3 (light blue) compared to the sample average (pre- and post-selection) in the sensitive wild-type parent is shown for each nuclear chromosome and the apicoplast genome. The figure was generated using the ggplots and tidyverse packages in Rstudio.

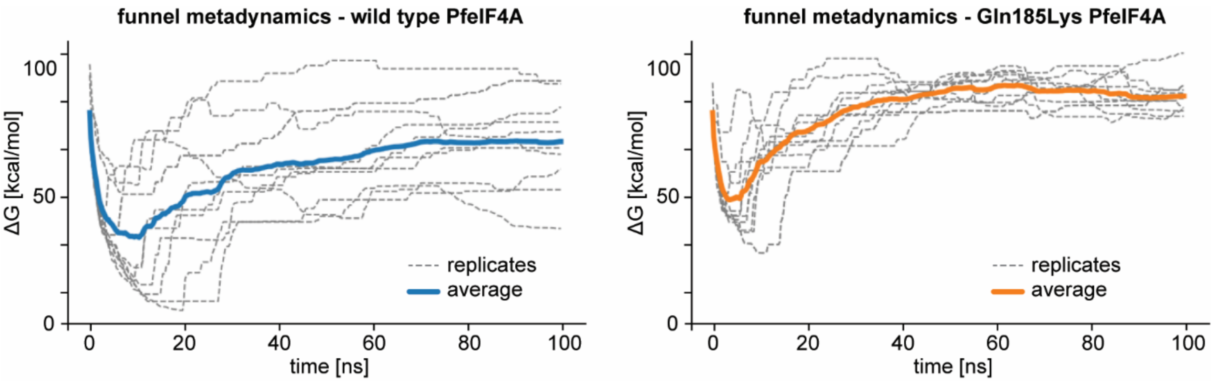

**Figure S3. Funnel metadynamics simulation.**

Simulations of protein-ligand interactions between CR-1-31-B and either wild type PfeIF4A (left panel) or the Gln185Lys mutant (right panel). Estimated binding free energy is indicated.

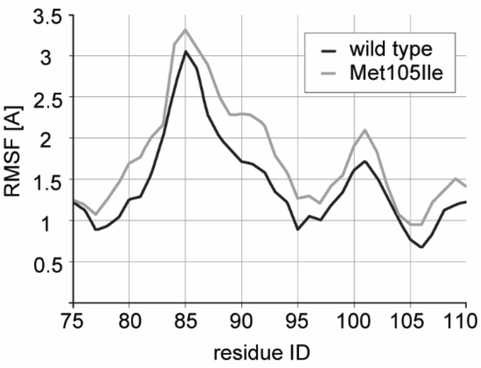

**Figure S4. Residue-wise root-mean-square fluctuations (RMSF) from the endpoint MD trajectories of the two end states used in the relative free energy calculations, comparing wild type Pf20S β5 (black line) and the Met105Ile mutant (grey line).**

The Met105Ile system shows consistently elevated flexibility relative to the wild type system, most prominently in the C-terminal region of the extended loop-β-hairpin-loop motif (residues 77–95) and in the segment comprising residues 105-110. These increased fluctuations are consistent with weakened stabilizing interactions in the mutant and support the observed reduction in hydrogen-bond occupancies involving Ser81, Gly107, Ala110, and Ala111, as well as the markedly diminished hydrogen-bonding frequency to the catalytic Thr1. Together, these changes provide a structural and dynamic rationale for the reduced probability of covalent reaction with the ONX-0914 warhead and the experimentally observed loss in binding affinity.

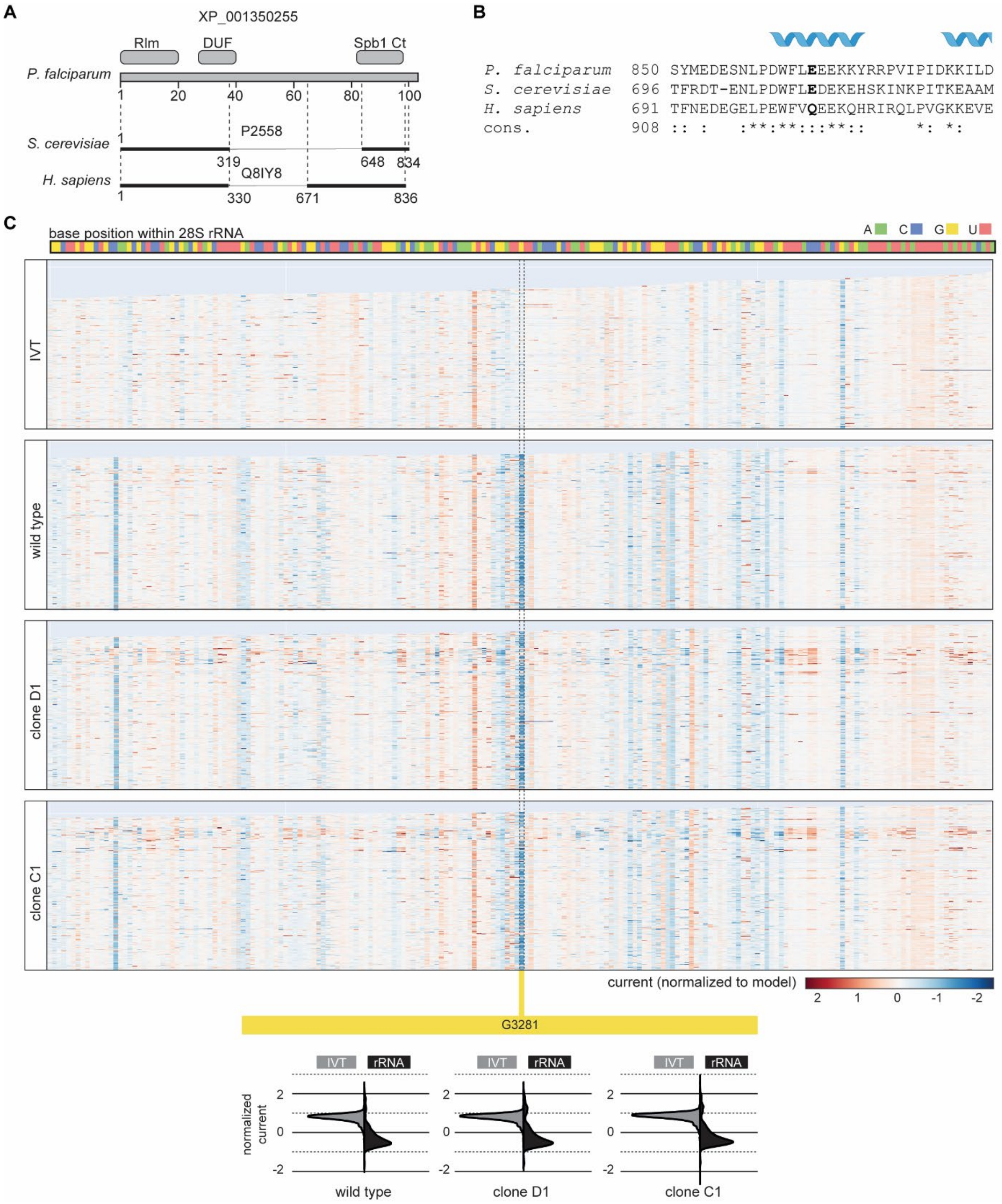

**Figure S5. The modification of rRNA base G3281 is conserved among eukaryotes and in *P. falciparum* likely mediated by putative PfRLM (PF3D7_1354300).**

**(A)** Graphical representation of a multiple sequence alignment of *P. falciparum* putative methyltransferase PfRLM (PF3D7_1354300), 27S pre-rRNA guanosine(2922)-2'-O)-methyltransferase SPB1 of *S. cerevisiae* (accession number P25582) and human pre-rRNA 2'-O-ribose RNA methyltransferase FTSJ3 (accession number Q8IY81). NCBI-annotated conserved domains are shown as boxes. RlmE, domain associated with members of the rRNA large subunit methyltransferase family E; DUF, domain of unknown function, Spb1 Ct, domain found at the C-terminus of SPB1 proteins. Conserved sequences are highlighted by black lines. **(B)** Amino acid sequence alignment of the region surrounding the mutation linked to brusatol-resistance (E864D) (highlighted in bold). The consensus (cons.) row indicates amino acids conserved in all three (*) or in two species (:). The alignment was performed with T-coffee (Di Tommaso *et al.*, 2011). Alphafold-predicted (Jumper *et al.*, 2021) alpha helices are shown in blue (confident prediction, 90 > pIDDT > 70). **(C)** Heatmaps showing Nanopore sequencing current at parasite 28S rRNA base positions of *in vitro* transcribed (IVT) and native rRNA isolated from the brusatol-sensitive parent (wild type) and resistant clones C1 and D1. Reads were aligned with Uncalled4 (Kovaka *et al.*, 2025). The color gradient represents the difference between the detected and expected normalized currents. rRNA base positions are indicated. Violin plots show the distribution of current measured using IVT rRNA bases (grey) and native parasite rRNA (black) at position G3281.

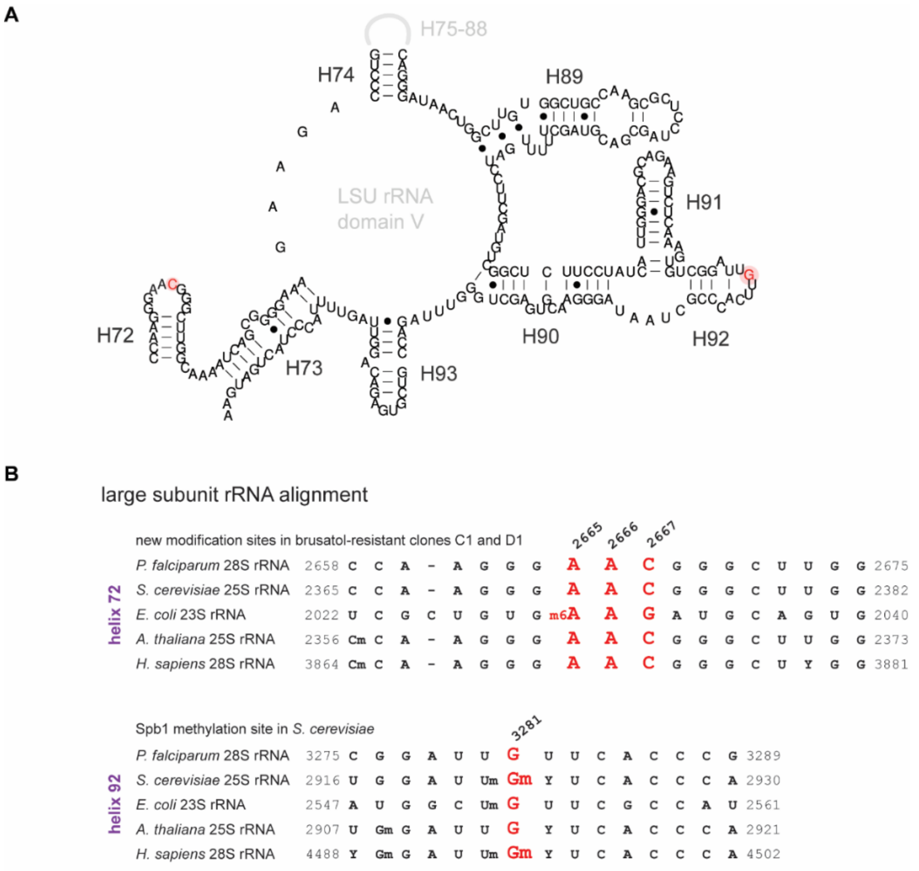

**Figure S6. The secondary structure and modification sites of eukaryotic rRNA surrounding the PTC.**
**(A)** Secondary structure segment of the *P. falciparum* 28S rRNA, containing parts of the rRNA involved in forming the PTC. Helices are numbered in dark grey. The following nucleotides are highlighted in red: G3281 in helix 92, which aligns to the ScSPB1-methylated Gm2922 in yeast; and C2667 exclusively 2'O-methylated in helix 72 of brusatol-resistant clones. These modifications are shown in the context of helix 89 – the presumed target of brusatol. The figure was created by aligning the *P. falciparum* 28S rRNA sequence to the resolved secondary structure of *Tetrahymena* 28S rRNA (Su *et al.*, 2021). **(B)** Alignment of rRNA sequences of helices 72 and 92 of the *P. falciparum* 28S rRNA and the corresponding sequences of the large rRNA subunit of *S. cerevisiae*, *E. coli*, *A. thaliana* and *H. sapiens* [sequences retrieved from RNAcentral (The RNAcentral Consortium *et al.*, 2026)]. Nm, base with 2'O-methylation; m$^6$A, N$^6$-methyladenosine. The modification status of the *S. cerevisiae, E. coli* and *H. sapiens* rRNA was obtained from Taoka *et al.*(Taoka *et al.*, 2018). The modification status of the *A. thaliana* rRNA was obtained from the modomics site (IIMCB), ID 229 (Sordyl *et al.*, 2026). As in (A), rRNA bases found at positions 3281 and 2667 in *P. falciparum* 28S rRNA are highlighted in red.

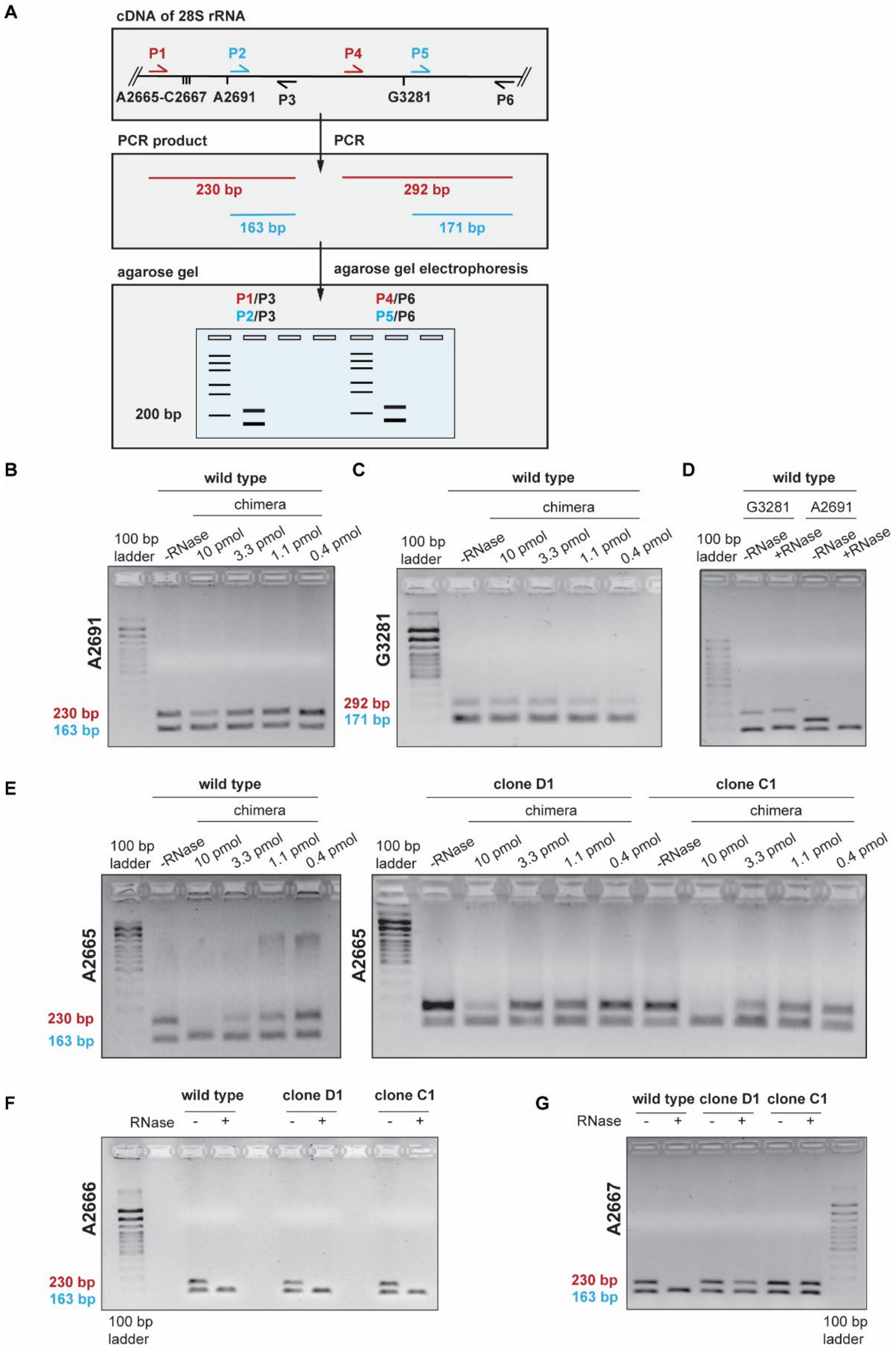

**Figure S7. Nm-VAQ detects 28S rRNA methylation status at base positions 2665-2667, 2691 and 3281 in brusatol-resistant parasites.**

**(A)** Schematic of the Nm-VAQ PCR primer design for identifying methylated 28S rRNA nucleotides (indicated as Nm). Two forward primers, one upstream and one downstream of the base of interest, and one reverse primer were used for PCR amplification of the cDNA template, resulting in a large (red) and short fragment (blue). If a 2'O-methylation is present at the suspected base, reverse transcription generates a large cDNA template (step not shown) that can be amplified by both primer pairs. If the base is 2'O-methylated, reverse transcription generates a short cDNA fragment that can only be amplified with the second primer pair (blue). **(B-D)** PCR products of cDNA derived from wild type 28S rRNA at the site surrounding (B, D) the "negative control" base 2691 and (C, D) the "positive control" base 3281 (n=2). **(E-G)** PCR products of cDNA derived from the wild type parent and brusatol-resistant clones C1 and D1 at the 28S rRNA sites surrounding base 2665 (E), 2666 (F) and 2667 (G).

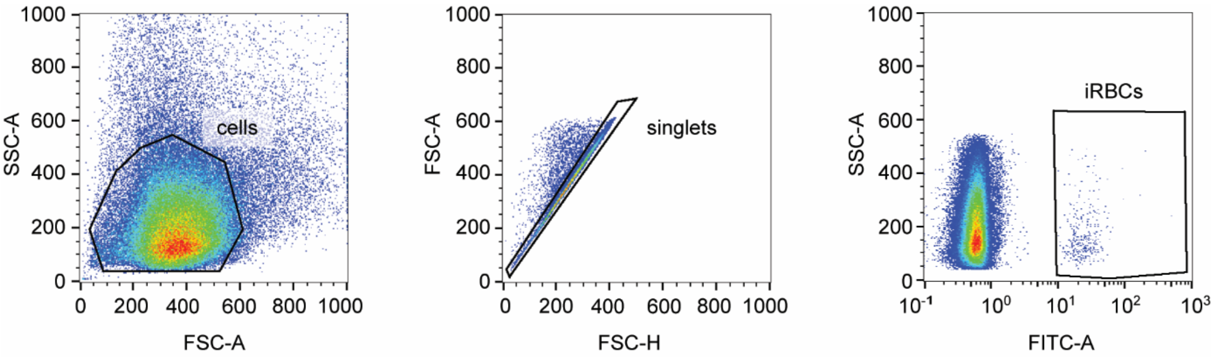

**Figure S8. Flow cytometry gating strategy.**
Representative flow cytometry plots of live SYBR Green I-stained asexual parasites are shown. Hierarchical gating was used to identify cells (SSC-A vs. FSC-A), singlets (FSC-A vs. FSC-H) and SYBR Green I-positive events (SSC-A vs. FITC-A). SSC, side scatter; FSC, forward scatter; FITC, fluorescein-5-isothiocyanate; iRBCs, infected red blood cells; A, area; H, height.

**Table S1. Parasites resistant to candidate drugs do not show cross-resistance.**

| parasite strain | resistance | CR-1-31-B $IC_{50}$ [nM] | brusatol $IC_{50}$ [nM] | ONX-0914 $IC_{50}$ [nM] |
|---|---|---|---|---|
| NF54 | | 1.065 ± 0.3551; n=3 | 7.821 ± 0.5860; n=2 | 16.16 ± 1.120; n=2 |
| RF12 | PPQ, CQ, PYR | 0.7878 ± 0.1126; n=4 | 6.755 ± 0.2040; n=2 | 13.59 ± 0.8350; n=2 |
| Dd2 | CQ, CYC, PYR | 0.8173 ± 0.1048; n=2 | 5.171 ± 0.4355; n=2 | 12.10 ± 1.065; n=2 |
| Dd2 (PfCARL$^{I1139K}$) | GNF156 | 0.7658 ± 0.0147; n=2 | 5.775 ± 0.4670; n=2 | 14.83 ± 0.0500; n=2 |
| Dd2 (PfATP4$^{G358S}$) | SJ557733 | 1.022 ± 0.0689; n=2 | 8.371 ± 0.0765; n=2 | 15.28 ± 0.6800; n=2 |
| Dd2 (PfACoAS$^{T648M}$) | MMV019721, MMV084978 | 0.8908 ± 0.1283; n=2 | 5.361 ± 0.8545; n=2 | 13.72 ± 0.1100; n=2 |
| Dd2 (PfPI4K$^{S743T}$) | MMV390048 | 0.7898 ± 0.0691; n=2 | 6.052 ± 0.9505; n=2 | 12.70 ± 0.4800; n=2 |
| Dd2 (PfDHODH$^{G181D}$) | DSM265 | 0.8558 ± 0.0138; n=2 | 6.358 ± 1.075; n=2 | 13.51 ± 1.050; n=2 |

$IC_{50}$ values for CR-1-31-B, brusatol and ONX-0914 determined in dose-response assays against asexual parasites of *P. falciparum* strains resistant to antimalarial drugs. The sensitive strain NF54 was used as a control. Values represent the mean ± s.e.m. of two to four biological replicate assays. PPQ, piperaquine; CQ, chloroquine; PYR, pyrimethamine, CYC, cycloguanil.

**Table S2. Resistance levels of parasite clones deriving from in vitro drug-selected populations.**

| compound | wild type parent | resistant clone | asexual $IC_{50}$ [nM] | $IC_{50}$ shift |
|---|---|---|---|---|
| CR-1-31-B | Dd2-B2 | | 1.26 ± 0.20 | 1.00 |
| | | A1 | 2.71 ± 0.19 | 2.15 |
| | | A2 | 3.51 ± 0.31 | 2.78 |
| | | A3 | 3.27 ± 0.19 | 2.59 |
| | | B1 | NA | >100 |
| | | B2 | NA | >100 |
| | | B3 | NA | >100 |
| brusatol | Dd2-B2 | | 7.04 ± 0.44 | 1.00 |
| | | C1 | 121.57 ± 21.13 | 17.26 |
| | | C2 | 106.22 ± 25.95 | 15.08 |
| | | C3 | 133.73 ± 40.64 | 18.99 |
| | | C4 | 130.77 ± 28.62 | 18.57 |
| | | D1 | NA | >100 |
| | | D2 | NA | >100 |
| | | D3 | NA | >100 |
| | | D4 | NA | >100 |
| | | D5 | NA | >100 |
| ONX-0914 | Dd2-B2 | | 16.541 ± 12.142 | 1.00 |
| | | E1 | 92.21 ± 2.91 | 5.74 |
| | | E2 | 91.38 ± 1.91 | 5.69 |
| | | E3 | 107.45 ± 2.38 | 6.69 |
| | | E4 | 97.60 ± 2.76 | 6.07 |
| | | E5 | 94.32 ± 2.36 | 5.87 |
| | | E6 | 103.16 ± 3.46 | 6.42 |
| | | E7 | 79.17 ± 1.54 | 4.93 |
| **compound** | **wild type parent** | **resistant clone** | **stage V $IC_{50}$ [nM]** | **$IC_{50}$ shift** |
| CR-1-31-B | Dd2-B2 | | 140.90 ± 13.07 | 1.00 |
| | | B1 | NA | >100 |
| brusatol | Dd2-B2 | | 119.25 ± 12.67 | 1.00 |
| | | C3 | 2612.52 ± 183.59 | 21.91 |
| | | D1 | NA | >100 |
| ONX-0914 | Dd2-B2 | | 29.72 ± 3.93 | 1.00 |
| | | E1 | 164.24 ± 30.96 | 5.53 |

$IC_{50}$ values of parasite clones resistant to CR-1-31-B, brusatol and ONX-0914. The sensitive Dd2-B2 parent served as a control. Dose-response assays were carried out using asexual parasites (top) and stage V gametocytes (bottom). Values represent the mean ± s.e.m. of three biological replicate assays. Dose-response assays of CR-1-31-B with asexual parasites were carried out in technical duplicates (s.e.m. is indicated). NA, not applicable.

**Table S3. SNPs identified in drug-resistant parasite clones.**

| **CR-1-31-B-resistant - population A and clones deriving from this population** | | |
|---|---|---|
| chromosome | Pf3D7_12_v3 | Pf3D7_14_v3 |
| position | 1620389 | 2825667 |
| geneID | PF3D7_1239000 | PF3D7_1468700 |
| reference | G | A |
| variant | A | G |
| format | GT:AD:DP:GQ:PL | GT:AD:DP:GQ:PL |
| parent_pre-selection | 0/0:144,0:144:99:0,120,1800 | 0/0:79,0:79:99:0,115,1800 |

| | | | |
|---|---|---|---|
| parent_post-selection | 0/0:51,0:51:99:0,120,1800 | 0/0:41,0:41:99:0,105,1037 | |
| clone A1 | 1/1:0,90:90:99:2635,270,0 | 1/1:0,68:68:99:1662,202,0 | |
| clone A2 | 1/1:0,106:106:99:3066,317,0 | 1/1:0,79:79:99:1916,235,0 | |
| clone A3 | 1/1:0,102:102:99:2983,306,0 | 1/1:0,73:73:99:1763,217,0 | |
| consequence of 1/1 | 3' UTR variant | Thr100Ala | |
| **CR-1-31-B-resistant - population B and clones deriving from this population** | | | |
| chromosome | Pf3D7_14_v3 | | |
| position | 2826044 | | |
| geneID | PF3D7_1468700 | | |
| reference | C | | |
| variant | A | | |
| format | GT:AD:DP:GQ:PL | | |
| parent_pre-selection | 0/0:107,0:107:99:0,105,1800 | | |
| parent_post-selection | 0/0:42,0:42:99:0,107,1101 | | |
| clone B1 | 1/1:0,111:111:99:3302,333,0 | | |
| clone B2 | 1/1:0,59:59:99:1753,177,0 | | |
| clone B3 | 1/1:0,102:102:99:3098,307,0 | | |
| consequence of 1/1 | Gln185Lys | | |
| **brusatol-resistant - population D and clones deriving from this population** | | | |
| chromosome | Pf3D7_11_v3 | Pf3D7_07_v3 | Pf3D7_13_v3 |
| position | 1510416 | 1103312 | 2164825 |
| geneID | PF3D7_1468700 | PF3D7_0726200 | PF3D7_1354300 |
| reference | G | A | T |
| variant | A | ATTATT | G |
| format | GT:AD:DP:GQ:PL | GT:AD:DP:GQ:PL | GT:AD:DP:GQ:PL |
| parent_pre-selection | 0/0:114,0:114:99:0,120,1800 | 0/0:79,0:79:99:0,119,1800 | 0/0:98,0:98:99:0,120,1800 |
| parent_post-selection | 0/0:51,0:51:99:0,120,1800 | 0/0:53,0:53:99:0,106,1318 | 0/0:46,0:46:99:0,106,1201 |
| clone D1 | 1/1:0,89:89:99:2668,267,0 | 1/1:0,60:60:99:2675,180,0 | 1/1:0,91:91:99:2284,272,0 |
| clone D2 | 1/1:0,112:112:99:3315,335,0 | 1/1:0,52:52:99:2301,156,0 | 1/1:1,93:94:99:2256,269,0 |
| clone D3 | 1/1:0,101:101:99:2999,302,0 | 1/1:0,58:58:99:2591,174,0 | 1/1:0,86:86:99:2114,256,0 |
| consequence of 1/1 | synonymous mutation | 3' UTR variant | Glu864Asp |
| **ONX-0914-resistant - population E and clones deriving from this population** | | | |
| chromosome | Pf3D7_10_v3 | | |
| position | 441641 | | |
| geneID | PF3D7_1011400 | | |
| reference | C | | |
| variant | T | | |
| format | GT:AD:DP:GQ:PL | | |
| parent_pre-selection | 0/0:100,0:100:99:0,104,1800 | | |
| parent_post-selection | 0/0:47,0:47:99:0,99,1257 | | |
| clone E1 | 1/1:0,86:86:99:2516,258,0 | | |
| clone E2 | 1/1:0,55:55:99:1619,165,0 | | |
| clone E3 | 1/1:0,122:122:99:3632,367,0 | | |
| consequence of 1/1 | Met105Ile | | |

Sequencing data is presented in the standard Variant Call Format (VCF) that includes information on alleles. As *P. falciparum* blood stage parasites are haploid, the GT (genotype) data shows either 0/0 (no difference to the reference) or 1/1 (different between the reference and drug-resistant clones). AD, allele depth (count of filtered reads supporting each allele, reference count/alternate count); GQ, genotype quality (phred-scaled confidence score that the assigned GT is correct); PL, phred-scaled likelihoods (normalized likelihoods for genotypes 0/0, 0/1, and 1/1).

**Table S4. rRNA base modifications.**

| | *E. coli* | | | *S. cerevisiae* | | | *H. sapiens* | | | *P. falciparum* | | |
|---|---|---|---|---|---|---|---|---|---|---|---|---|
| **alignment** | position | base | modification | position | base | modification | position | base | modification | position | base | modification |
| 4281 | 2029 | G | - | 2369 | G | - | 3870 | G | - | 2664 | G | - |
| 4282 | 2030 | A | $m^6A$ | 2370 | A | - | 3871 | A | - | 2665 | A | - |
| 4283 | 2031 | A | - | 2371 | A | - | 3872 | A | - | 2666 | A | - |
| 4284 | 2032 | G | - | 2372 | C | - | 3873 | C | - | 2667 | C | - |
| 4973 | 2913 | G | - | 2920 | G | Gm | 4494 | G | Gm | 3281 | G | Gm |
| 4871 | 2451 | A | - | 2818 | A | - | 4392 | A | - | 3179 | A | - |
| 4872 | 2452 | C | - | 2819 | C | - | 4393 | C | - | 3180 | C | - |
| 4484 | 2201 | G | - | 2509 | A | - | 4068 | A | - | 2820 | A | - |
| 4485 | 2202 | U | - | 2510 | C | - | 4069 | C | - | 2821 | C | - |

Post-transcriptional modifications of selected rRNA bases of the large ribosomal subunits. Nm, base with 2'O-methylation; $m^6A$, $N^6$-methyladenosine.

**Table S5. Oligonucleotides used in this study.**

| **plasmid cloning** | | | |
|---|---|---|---|
| **plasmid** | **oligonucleotide** | **sequence 5' to 3'** | **fragment name** |
| pHF_gC-cg6#3 | cg6#3_F | **tatt**ctttaaaaatttattcgaac | sgt_*cg6#3* |
| | cg6#3_R | **aaac**gttcgaataaatttttaaag | |
| pD_cg6_pfs16-RE9H | gib1_F | *cgttggccgattcattaatg*AGGTGTTGCTCAAATAGTGTCG | PCR fragment 1 (5' box) |
| | gib4_R | *ggatggatatttatttttggatagctac*GTTCGAATAAATTTTTAAAGCATTTC | |
| | gib3_F | *gaaatgctttaaaaatttattcgaac*GTAGCTATCCAAAAATAAATATCCATCC | PCR fragment 2 (*pfs16* promoter) |
| | gib6_R | *gatgttcttggcgtcctccat*GTTGAAGAAAGTATAAATAGAAAAATGGC | |
| | gib5_F | *gccatttttctatttatactttcttcaac*ATGGAGGACGCCAAGAACATC | PCR fragment 3 (*re9h* cds) |
| | gib8_R | *ctattattaaataaat*TCAGATCTTGCCGCCCTTCTTGGC | |
| | gib7_F | *gccaagaagggcggcaagatctga*ATTTATTTAATAATAGATTAAAAATATTATAAAAATAAAAAC | PCR fragment 4 (*hrp2* terminator) |
| | gib10_R | *catgcaattcttgcaacttgtctatg*TTTAATAAATATGTTCTTATATATAATGAG | |
| | gib9_F | *ctcattatatataagaacatatttattaaa*catagacaagttgcaagaattgc | PCR fragment 4 (3' box) |
| | gib12_R | *cctcttcgctattacgccag*aaacatatcatatggaaactattccc | |
| | gib11_F | *gggaatagtttccatatgatatgttt*CTGGCGTAATAGCGAAGAGG | PCR fragment 6 (pD backbone) |
| | gib2_R | *cgacactatttgagcaacacct*CATTAATGAATCGGCCAACG | |

| Primer binding sites are highlighted in capital letters. Sequences involved in Gibson assemblies are highlighted in italics. Single-stranded overhangs required for T4 DNA ligase-dependent cloning of the double-stranded sgRNA target sequence are highlighted in bold. F, forward; R, reverse | |
|---|---|
| **diagnostic PCRs on genomic DNA** | |
| **oligonucleotide** | **sequence 5’ to 3’** |
| p1 | aagctctcgggcacgaagt |
| p2 | aagaccctgggcgtgaacca |
| p3 | tgatactaatcaaactgatgaggaag |
| p4 | agcaggtaaatgagcaagg |
| p5 | acatataattgaggaaatgtaggcc |
| p6 | actggcctacatttcctc |

| **Nm-VAQ oligonucleotides** | | |
|---|---|---|
| **Site** | **oligonucleotide** | **sequence 5’ to 3’** |
| 2665 | chimera | aaaccacagccaaggga |
| | RT primer 1 | aatatcagaatgccttcccg |
| | reverse primer | gccttcccgaaggataaagc |
| | forward primer upstream | ctatctagcgaaaccacagc |
| | forward primer downstream | agctttactctagtctggc |
| 2691 | chimera | gcaaaatcagcggggaa |
| | RT primer 2 | aatatcagaatgccttcccg |
| | reverse primer | gccttcccgaaggataaagc |
| | forward primer upstream | ctatctagcgaaaccacagc |
| | forward primer downstream | agctttactctagtctggc |
| 3281 | chimera | ctcaaagtgtcggattg |
| | RT primer | gtcaaccaattgctgtacc |
| | reverse primer | gtcaaccaattgctgtaccaattgtc |
| | forward primer upstream | ttaccacagggataactggc |
| | forward primer downstream | cccgctaatagggaacgtgagc |

| **IVT PCR primers** | |
|---|---|
| **oligonucleotide** | **sequence 5’ to 3’** |
| A2_F1 | taatacgactcactataggggtttgatttgatgttaggagggcaaatc |
| A2_R1 | ttttttttttttttttttttttttttttttccatctaagaaactcataatactattggcacg |
| A2_F2 | taatacgactcactatagggggttccttccccgtctcaaaagg |
| A2_R2 | ttttttttttttttttttttttttttttttaaacactttaccactagggataagcctc |

**Dataset S1. Results from the high-throughput primary and validation screens of 51,185 compounds against mature stage V gametocytes.**

Primary screen (worksheet 1): Activity values (% inhibition of gametocyte viability) for all 51,185 compounds screened at 5 µM against mature stage V gametocytes (day 12). Values represent the reduction in relative luminescence counts normalized to the mean luminescence signals from negative control wells (untreated, 0% activity) and positive control wells (50 µM MB, -100% activity). MoA, mode-of-action library; NAPR, natural product library.
Validation screen (worksheet 2): $IC_{50}$ values, % activity at maximum concentration ($C_{max}$) and selectivity indices (SI) determined from 8-point dose-response assays against mature stage V gametocytes (day 12) and HepG2 cells for 844 compounds that showed > 30% inhibition of gametocyte viability in the primary screen. For 156 compounds with gametocytocidal $IC_{50}$ < 2 µM, dose-response assays have also been performed on asexual parasites. Results from structural clustering of all compounds with gametocytocidal $IC_{50}$ < 2 µM are shown in column ‘cpd cluster’.

Final hit list (worksheet 3): All compounds with gametocytocidal $IC_{50}$ < 0.5 µM and SI > 10, including structures. Hits analyzed further in this study are highlighted in yellow. For compounds 390 and 464, order numbers (Mcule, Inc.) are given for reference.

**Dataset S2. Copy number variation analysis.** Gene copy number (CN) for wild type population pre-selection (wt1) and at the end (wt2) of resistance selection, brusatol-resistant clones D1, D2 and D3. The suffix following the gene identifier denotes the specific architectural components of the gene feature: **.1** identifies the transcript or splice isoform number, **-p1** indicates the primary translated peptide product, and **-CDS1** specifies the first continuous protein-coding sequence (exon segment). Chromosomal positions of beginning (start) and end (stop) of the architectural components are listed. CN was determined by dividing the gene-specific reads per kilobase per million (RPKM) by the average RPKM. CN ratios between each sample and the wt1 sample are color-coded.
